# Pervasive conservation of intron number and other genetic elements revealed by a chromosome-level genomic assembly of the hyper-polymorphic nematode *Caenorhabditis brenneri*

**DOI:** 10.1101/2024.06.25.600681

**Authors:** Anastasia A. Teterina, John H. Willis, Charles F. Baer, Patrick C. Phillips

## Abstract

With within-species genetic diversity estimates that span the gambit of that seen across the entirety of animals, the *Caenorhabditis* genus of nematodes holds unique potential to provide insights into how population size and reproductive strategies influence gene and genome organization and evolution. Our study focuses on *Caenorhabditis brenneri*, currently known as one of the most genetically diverse nematodes within its genus and metazoan phyla. Here, we present a high-quality gapless genome assembly and annotation for *C. brenneri*, revealing a common nematode chromosome arrangement characterized by gene-dense central regions and repeat rich peripheral parts. Comparison of *C. brenneri* with other nematodes from the ‘Elegans’ group revealed conserved macrosynteny but a lack of microsynteny, characterized by frequent rearrangements and low correlation iof orthogroup sizes, indicative of high rates of gene turnover. We also assessed genome organization within corresponding syntenic blocks in selfing and outcrossing species, affirming that selfing species predominantly experience loss of both genes and intergenic DNA. Comparison of gene structures revealed strikingly small number of shared introns across species, yet consistent distributions of intron number and length, regardless of population size or reproductive mode, suggesting that their evolutionary dynamics are primarily reflective of functional constraints. Our study provides valuable insights into genome evolution and expands the nematode genome resources with the highly genetically diverse *C. brenneri*, facilitating research into various aspects of nematode biology and evolutionary processes.

## Introduction

A central tenet of molecular evolution is that the pace and structure of the evolution of genomes and genetic elements should be largely determined by variation in population size. This is because it is thought that the expansion of some genetic elements, such as repetitive DNA and introns, should be weakly deleterious (Burns 2022; Carmel and Chorev 2012), and therefore be expected to grow in small populations (4*N*_e_*s* < 1, where *N*_e_ is the effective population size and s is the selection coefficient) and be eliminated in large populations (4*N*_e_*s* > 1) through a balance of natural selection and genetic drift. For example, Lynch and colleagues have suggested that major determinant of differences in genome and gene structures among prokaryotes and eukaryotes is due to differences in effective population size (Lynch 2002; Lynch 2007, however, see discrepancies in Daubin and Moran 2004; Vinogradov 2004; Charlesworth and Barton 2004). Within animals there tends to be some signature of the influence of population size on genome size and structure (Gregory et al. 2007; Galtier 2024), but the results are often equivocal (Roddy et al. 2021; Marino et al. 2024). Often vast evolutionary distances are included in these comparisons, which can lead to challenges of phylogenetic non-independence in the analyses (Whitney and Garland 2010; Whitney et al. 2011), as well as the likelihood that the actual functional role of genetic elements of interest may themselves have changed. Nematodes of the family *Caenorhabditis* hold the promise to circumvent some of these challenges, as species within this group include some of the least diverse and most diverse animals known (Noble et al. 2021; Dey et al. 2013), indicating vast differences in effective population size. Until recently, however, complete genomes have only been available for a few selfing species, including the well-studied model system *C. elegans* (The C. elegans Sequencing Consortium 1998; Stein et al. 2003; Noble et al. 2021). Genome assemblies of outcrossing species have frequently been contaminated by residual polymorphism, although this has been addressed using long-read sequencing in a several species with moderate to high levels of within-species polymorphism (Barrière et al. 2009; Kanzaki et al. 2018; Teterina et al. 2020). Here, we push the limits of this comparative framework by presenting a whole-chromosome assembly of one of the most polymorphic animals currently known (Dey et al. 2013; Alm Rosenblad et al. 2021), *C. brenneri*, and use this new assembly in a comparative analysis of the evolution of size and structure of a variety of genetic elements thought to be subject to nearly-neutral evolution, particularly introns.

*Caenorhabditis* nematodes have compact genomes, typically 80-140Mb, with six holocentric chromosomes — five autosomes and one sex chromosome (Brenner 1974; Pires-daSilva 2007; Sun et al. 2022). While the majority of *Caenorhabditis* species have males and females (dioecy) and reproduce via outcrossing, three species have independently shifted to self-reproduction, or “selfing,” with most individuals being hermaphrodites with a few males (androdioecy, Kiontke et al. 2011). This mode of reproduction has greatly reduced the effective population size of these species (Golding and Strobeck 1980; Pollak 1987; Nordborg and Donnelly 1997; Thomas et al. 2012; Cutter 2019). Genomes of outcrossing species are larger and contain a greater number of genes than selfing species (Bird et al. 2005; Fierst et al. 2015; Yin et al. 2018; Stevens et al. 2019; Adams et al. 2023), with some of the reduction being attributable to the shift to self-reproduction per se via resulting changes in the regulation of male-specific pathways (Thomas et al. 2012; Yin et al. 2018; Sánchez-Ramírez et al. 2021;. Xie et al. 2022). Nematodes have a conservative chromosome organization, with high gene density in the central parts of chromosomes and repetitive elements concentrated in peripheral “arms” (The C. elegans Sequencing Consortium 1998; Carlton et al. 2022). Additionally, they exhibit conservative macro-synteny across Rhabditid species, with most orthologous genes consistently located on homologous chromosomes (De La Rosa et al. 2021; Tandonnet et al. 2019). Despite this, *Caenorhabditis* nematodes display a relatively high rate of gene turnover, involving both the expansion and reduction of gene families (Fierst et al. 2015; Adams et al. 2023; Ma et al. 2024). Moreover, the exon/intron structure in orthologous genes in nematodes is not conservative, particularly via the substantial loss of introns (Kent and Zahler 2000; Cho et al. 2004; Kiontke et al. 2004; Coghlan and Wolfe 2004; Ma et al. 2022).

*Caenorhabditis* nematodes are especially useful for testing hypotheses regarding the impact of population size on gene and genome organization because of the very substantial differences in population sizes across species. Namely, outcrossing nematode *C. brenneri*, named in honor of Sydney Brenner for his pivotal role in *C. elegans* research, has an effective population size of about 10^7^, making its genetic diversity on the order of many bacterial, with nearly one in ten nucleotides being polymorphic (Sudhaus and Kiontke 2007; Dey et al. 2013). Another outcrossing nematode *C. remanei* displays an effective population size of around 10^6^ (Graustein et al. 2002; Cutter et al. 2006; Teterina et al. 2023), whereas selfing species like *C. elegans*, *C. briggsae*, and *C. tropicalis* exhibit smaller population sizes, approximately 10^4^ (based on diversity estimations from Sivasundar and Hey 2003; Barrière and Félix 2005; Cutter 2006; Dolgin et al. 2007; Dolgin et al. 2008; Noble et al. 2021 considering the mutation rate from Saxena et al. 2019). And these differences translate into substantial differences in the probability of fixation of deleterious variation in introns (fig. 1, Lynch 2002 c.f. Eq. 1a). If assumptions underlying these expected outcomes hold, then we would predict that we should see noticeable disparities in intron size distributions across nematode species. To effectuate the test of this predication, we generated a gapless chromosome-scale genome assembly and high-quality annotation for *C. brenneri* and assessed divergence and conservation of the genomic landscape of genetic elements across the genus using a comparative framework. As might be anticipated, no single simple factor explains the overall patterns of genomic evolution that we observe, which are characterized by macro-level conservation and micro-level evolutionary dynamics.

**Fig. 1.**
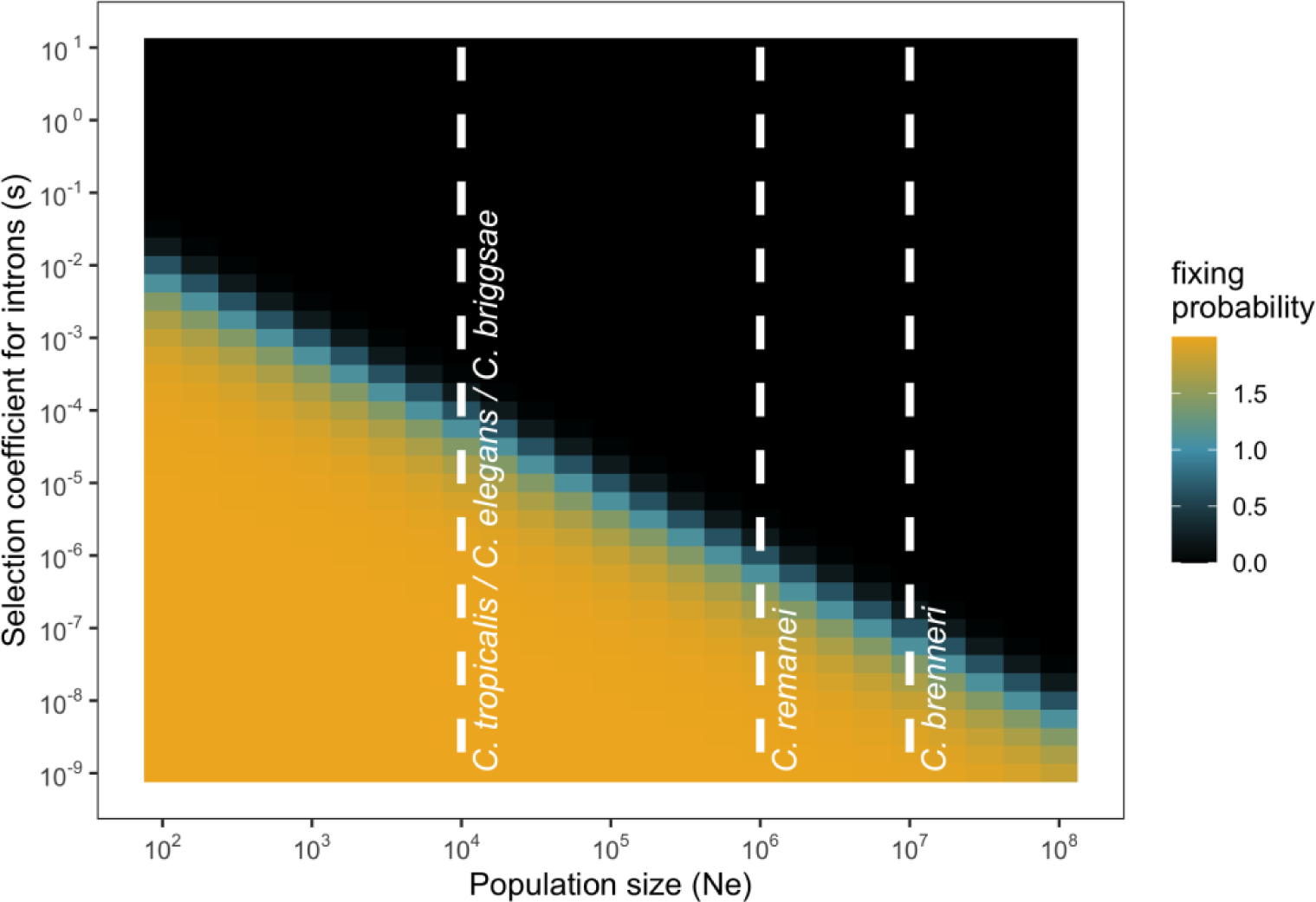
Scaled probabilities of fixation of introns in diploid populations. The values are based on Equation 1a from Lynch (2002). The white dashed lines demonstrate the order of population sizes for some *Caenorhabditis* nematodes.

## Results

### Chromosome-scale genome assembly of Caenorhabditis brenneri

Using 90 generations of brother-sister mating, we generated a highly inbred strain CFB2252 of *C. brenneri* with greatly reduced residual heterozygosity. Using a combination of long- and short-reads, Hi-C data, and high-accuracy long-read (HiFi) data from an individual nematode, we assembled and scaffolded a gapless genome for *C. brenneri*. Interestingly, the genome exhibited substantial continuity following the initial assembly with long CLR PacBio reads (supplementary fig. S1, Supplementary Material), likely due to the low level of genetic variation in the CFB2252 strain (residual k-mer based heterozygosity of 0.8%), with the entire genome being fully assembled with the inclusion of Hi-C data (supplementary fig. S1, Supplementary Material). Also, we used HiFi reads to generate a mitochondrial genome (supplementary fig. S2, Supplementary Material), which matches the previously available mitochondrial genome (NC_035244.1). After removal of the assembly artifacts and bacterial contamination, we were left with 163 alternative haplotypes that were added to the assembly as unplaced scaffolds. Our new *C. brenneri* assembly shows tremendous improvement of quality and reduction on the duplication level in BUSCO genes from ∼30% to 1% as compared to the previously available assembly (Table 1).

**Table 1.** Assembly statistics for *C. brenneri* assemblies. Benchmarking Universal Single-Copy Orthologs (BUSCO) statistics were estimated for the databases version 10 using all scaffolds.

|  |  |  |
| --- | --- | --- |
| *C. brenneri* genome assembly | This study | The *Caenorhabditis brenneri* Sequencing and Analysis Consortium |
| Accession number | GCA_964036135.1 | GCA_000143925.2 |
| Strain | CFB2252 | PB2801 |
| Total ungapped length, bp | 123,187,963 | 170,093,652 |
| Percent gaps,% | 0 | 10 |
| Number of scaffolds | 6 chromosomes + mtDNA (+ 163 alternative haplotypes) | 3,305 |
| Scaffold N50 | 20 Mb | 382 Kb |
| Scaffold L50 | 3 | 120 |
| GC% | 38.42 | 38.5 |
| QV | 36 | - |
| Number of protein-coding genes | 25,756 | 30,670 |
| BUSCO Metazoa, 954 genes | Completeness:71.8%<br>[Single-copy:70.6%, Duplicated:1.2%]<br>Fragmented:3.7%<br>Missing:24.5% | Completeness:72.5%<br>[Single-copy:45.6%, Duplicated:26.9%]<br>Fragmented:3.5%<br>Missing:24.0% |
| BUSCO Nematoda, 3131 genes | Completeness:97.3%<br>[Single-copy:96.6%, Duplicated:0.7%]<br>Fragmented:1.1%<br>Missing:1.6% | Completeness:97.2%<br>[Single-copy:65.4%, Duplicated:31.8%]<br>Fragmented:1.0%<br>Missing:1.8% |

The genome organization of *C. brenneri* exhibits a domain-like structure similar to several other *Caenorhabditi*s species (The C. elegans Sequencing Consortium 1998; Carlton et al. 2022; Woodruff and Teterina 2020) with more repeats and lower gene density in the peripheral arms of the chromosomes and greater gene density in central regions across all chromosomes (fig. 2). Notably, *C. brenneri* shows a relatively low percentage of repeats (only 15.9%), which is lower than other species of *Caenorhabditis* (Fierst et al. 2015; Woodruff and Teterina 2020). This difference may be attributed to the divergence of the repeats from one another, as the current repeat annotation protocols only mask repeats that are 80% identical.

**Fig. 2.**
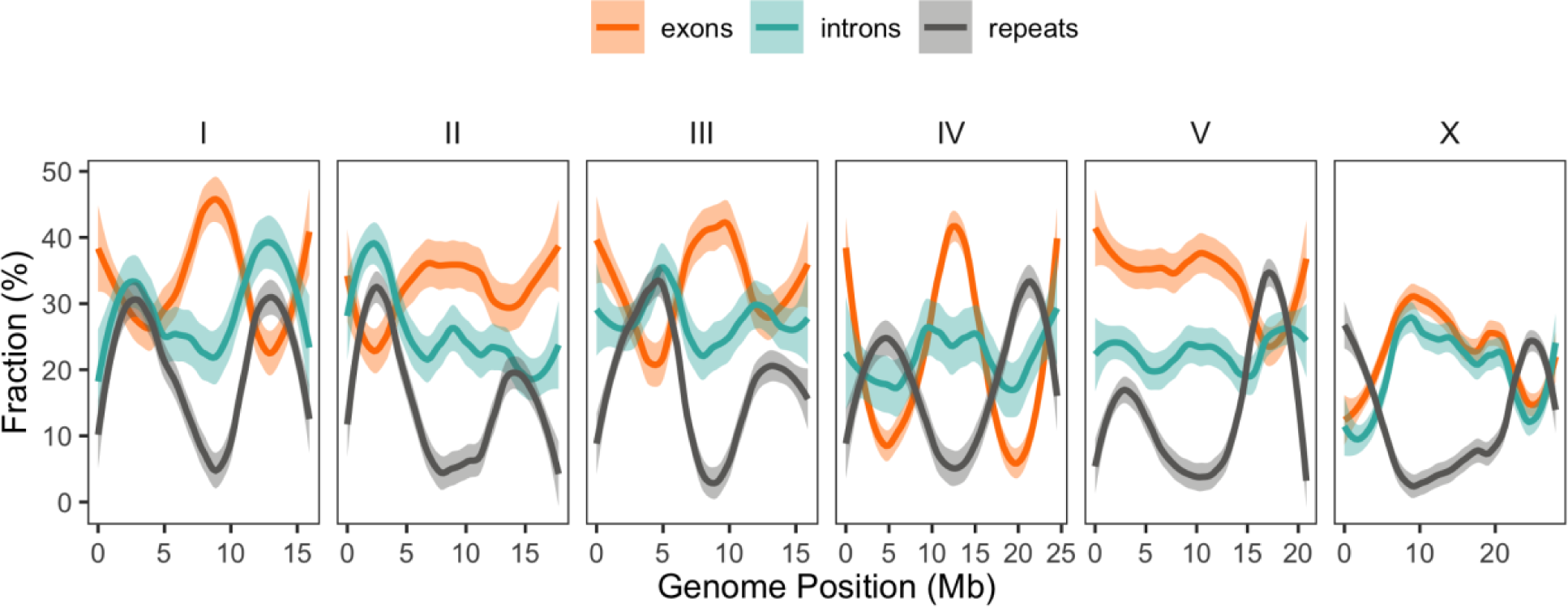
Genomic landscape of genetic elements for *C. brenneri*. Colored lines represent the smoothed means of the fraction of repetitive DNA (gray), exons (orange), and introns (teal) calculated from 100-kb windows. Shaded areas show 95% C.I.s of the mean.

### Central domains of chromosomes are more conservative

To examine the conservation pattern along the genome, we aligned the *C. brenneri* genome with genomes of several closely related *Caenorhabditis* nematodes. The phylogenetic trees, built from 50kb windows alignments, supports a sister-species relationship of *C. brenneri* with *C. sp48* throughout the genome (fig. 3A and B). Only a few windows (2.1% of the total) displayed a different tree topology; however, they were primarily located in the most divergent regions where reconstruction is likely less reliable. The tree heights, as well as the lengths of *C. brenneri* and *C. sp48* clade branches, were longer in the peripheral parts of chromosomes (two top rows in fig. 3C), indicating greater conservation in the central regions of chromosomes. This observation in agreement with previous findings in nematodes (The C. elegans Sequencing Consortium 1998; Hillier et al. 2007; Teterina et al. 2020). Additionally, we calculated the mean conservation score for each 50-kb window alignment using the phastCons algorithm (Siepel et al. 2005), a phylogenetic hidden Markov model-based method that estimates the probability of each nucleotide being under negative selection and belonging to conservative elements based on sequence alignments. The central regions exhibited elevated conservation scores (fig. 3C, bottom row), consistent with the distribution of tree heights. This pattern of central-domain conservation is likely associated with the high gene density as well as the presence of genes with more conserved function being located in the central parts of chromosomes (Parkinson et al. 2004; Cutter et al. 2009). Further, suppressed recombination in the central domain that has been observed in several other closely related species (Rockman and Kruglyak 2009; Ross et al. 2011; Teterina et al. 2023). When combined with natural selection, these factors can drive down local diversity and divergence through the action of linked selection (Smith and Haigh 1974; Charlesworth et al. 1993; Nordborg et al. 1996; Coop and Ralph 2012).

**Fig. 3.**
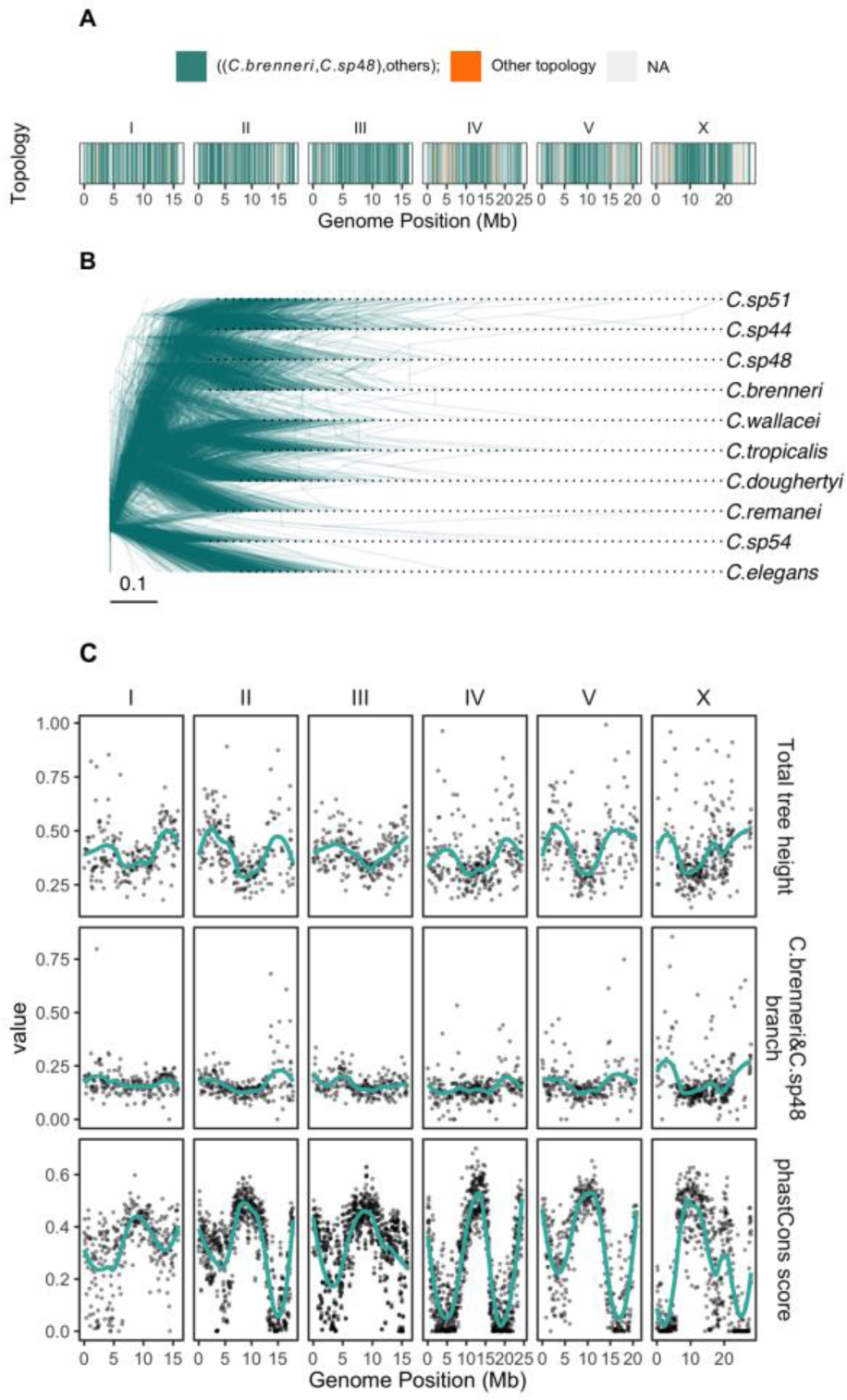
Divergence landscape and phylogeny of *C. brenneri*. **(A)** The tree topology along the genome of *C. brenneri* in 50-Kb windows. Teal color indicates that *C. brenneri* is the sister species with *C. sp48*, while orange represents other topologies, and light grey indicates missing data. **(B)** Maximum likelihood trees sampled across the genome (500 trees), derived from 50 kb non-overlapping windows in whole-genome alignments. **(C)** Tree statistics along the genome of *C. brenneri*. The top row displays total tree heights; the middle row illustrates the length of *C. brenneri* and *C. sp48* branch or *C. brenneri* branch for trees where they are not sister species. Conservation scores along the genome, estimated using the phastCons algorithm, are shown in the bottom row.

To examine the pattern of chromosome rearrangements and the evolution of genome organization, we utilized orthogroups and conservative gene order (Lovell et al. 2022) to infer syntenic blocks across *C. brenneri* and six other *Caenorhabditis* species with chromosome-scale assemblies (fig. 4). *Caenorhabditis* nematodes have six holocentric chromosomes that show almost no interchromosomal rearrangements but exhibit a significant number of intrachromosomal translocations and inversions, except for *C. nigoni* and *C. briggsae*, closely related species capable hybridizing with one another (Woodruff et al. 2010), these nematodes displayed only a few rearrangements. Statistics on the number and sizes of syntenic blocks relative to *C. brenneri* are provided in Table S1. Naturally, *C. brenneri* had the smallest number and longest syntenic blocks with the most closely related species, *C. tropicalis*. The number of blocks in reversed orientation was noticeably high, ∼50%, however, the mean lengths of blocks in reverse orientation tend to be smaller. Repression of interchromosomal rearrangements in nematodes and the stability of karyotypes have been described in other work (Stein et al. 2003; Hillier et al. 2007; Kanzaki et al. 2018; Teterina et al. 2020; Stevens et al. 2020; Sun et al. 2022; De La Rosa et al. 2021) and could potentially be a consequence of large population sizes and selection against large interchromosomal translocations, meiotic control mechanism such as pairing centers (McKim et al. 1993; Villeneuve 1994; reviewed in Carlton et al. 2022), and/or other mechanisms controlling genome stability during meiosis (Hillers and Villeneuve 2003; Lui and Colaiácovo 2013; Bhargava et al. 2020; Dokshin et al. 2020).

**Fig. 4.**
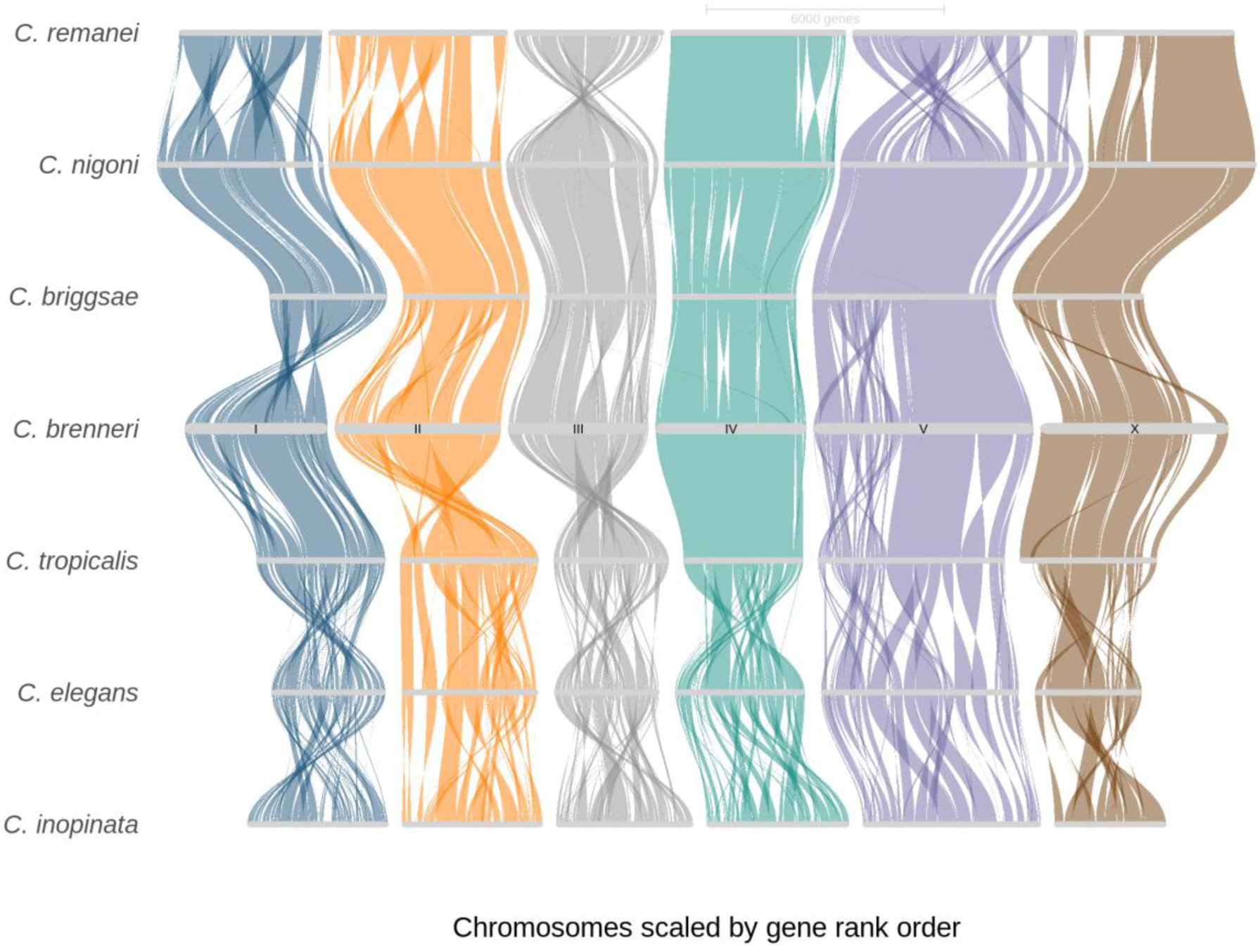
The riparian plot shows syntenic blocks among *Caenorhabditis* nematodes, based on the order of orthologues genes, demonstrating substantial intrachromosomal rearrangements. The central domains exhibit more extended synthetic blocks. *C. brenneri* was used as a reference. Also, chromosome IV of *C. remanei* was rotated.

### Genome organization in selfing and outcrossing species

Self-reproduction can affect genomic organization due to higher genetic drift relative to outcrossing species. It has been demonstrated that selfing nematodes have 10-30% smaller genomes than outcrossing ones, predominately due to the gene loss (Fierst et al. 2015; Yin et al. 2018; Adams et al. 2023). Subsequent to these studies, additional chromosome-scale genomes for outcrossing species of the Elegans group have become available (Kanzaki et al. 2018; Teterina et al. 2020; and this work), enabling us to investigate specific features of genome organization that distinct selfing from outcrossing species. We employed both generalized linear models and gradient boosting, a machine learning method based on ensemble learning with decision trees, to classify species as either outcrossing or selfing. The classification criteria included ratios of fractions and numbers of exons, introns, and genes, as well as the size of the blocks and intergenic regions within a species, relative to the outcrossing species *C. brenneri* (supplementary fig. S3, Supplementary material). These comparisons were made within inferred syntenic blocks, which were replicated proportionally to their size in *C. brenneri*. The models were trained on two selfing species, *C. elegans* and *C. tropicalis*, and two outcrossing species, *C. inopinata* and *C. remanei*. Subsequently, we tested the models on ratios from two closely related species, selfing *C. briggsae* and *C. nigoni*. The results from the generalized linear model were more accurate than those from the gradient boosting model approach (supplementary fig. S4, Supplementary material). We were able to successfully classify selfing and outcrossing species. The fraction of intergenic DNA and exon count were the most important features for classification (fig. 5A). Fig. 5B shows the coefficients and significance for various factors and their interactions used in the generalized linear model, and fig. 5C displays the permutation importance of those factors. The results support prior findings, demonstrating that selfing species lose a considerable number of genes but also a substantial amount of intergenic non-coding DNA (Fierst et al. 2015; Yin et al. 2018; Cutter et al. 2019; Adams et al. 2023). More extensive annotation of various functional and regulatory elements should allow these discrepancies to be explored in greater detail.

**Fig. 5.**
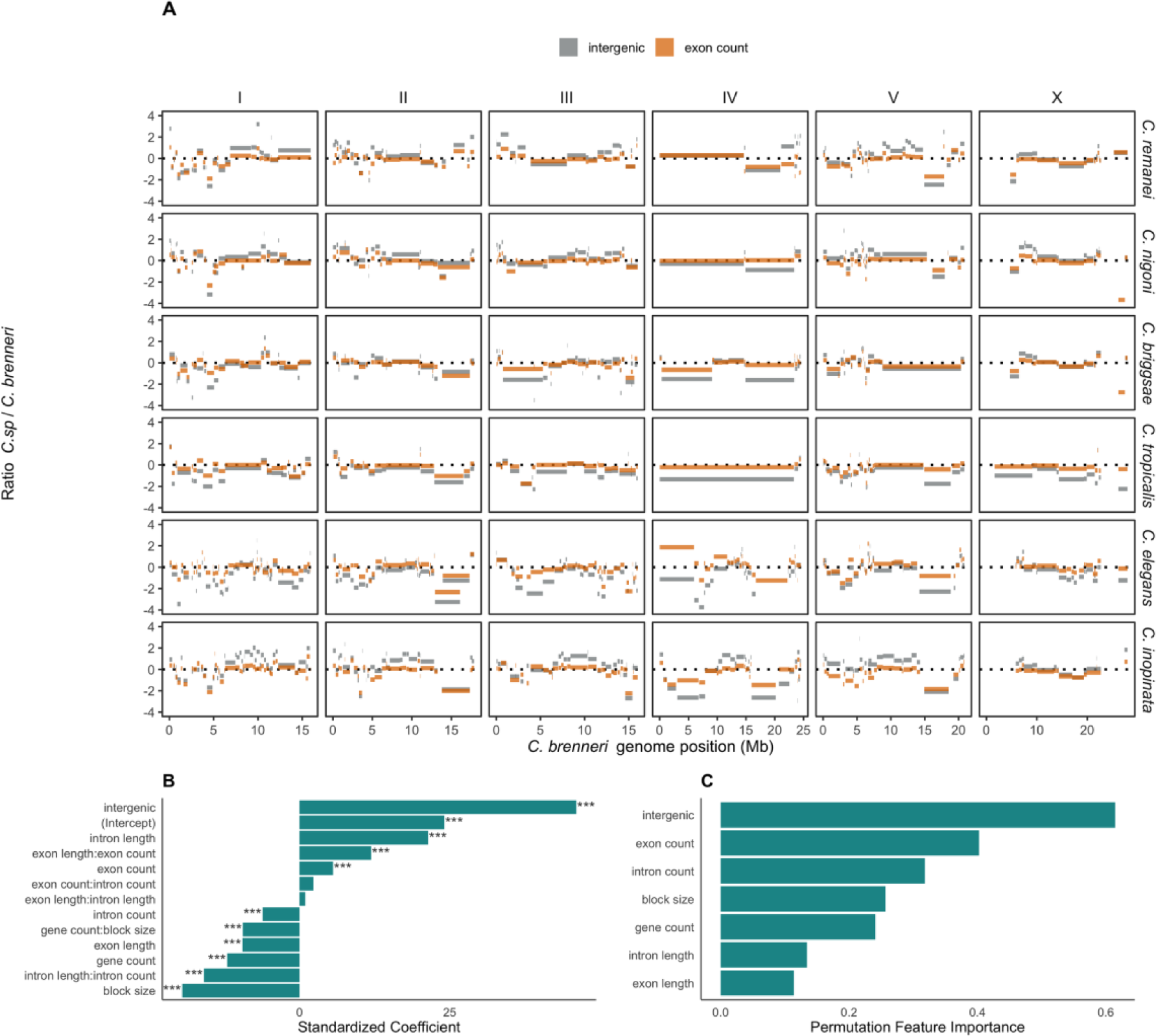
Comparison of genetic elements in *Caenorhabditis* species relative to *C. brenneri* and classification of selfing and outcrossing species using those features. **(A)** Ratios of exon counts (orange) and fraction of intergenic regions (grey) between a focal species and *C. brenneri* in corresponding syntenic blocks (see fig. 4) along the *C. brenneri* genome. **(B)** Coefficients of genomic features used in the generalized linear model for classifying selfing and outcrossing species. *** denotes Bonferroni-corrected p-values < 0.0001. **(C)** Permutation importance of features for classifying nematodes into selfing and outcrossing species.

### Influence of population size on gene organization

To explore the exon/intron structure of genes in nematodes, we compared orthologous genes among seven *Caenorhabditis* species with chromosome-scale assemblies. Nematodes display a low correlation in orthogroup sizes that rapidly decays with phylogenetic distance (fig. 6A). This observation is in accordance with data indicating high rates of gene turnover (Adams et al. 2023; Ma et al. 2024). However, when estimating the difference in sizes of shared orthogroups, the correlations were close to zero (supplementary fig. S5A, Supplementary Material), and the percentage of shared orthogroups varies among species pairs. Supplementary fig. S5B (Supplementary material) illustrates the percentage of shared orthogroups and the total number of genes in them. Outcrossing nematodes have on average 2.4 times more species-specific orthogroups than selfing species. We subsequently aligned exon/intron structures of 7,238 single-copy (hereinafter, 1-to-1) orthogroups. Several examples of such alignments are demonstrated in supplementary fig. S6 (Supplementary material). There, the majority of coding sequences are conserved and present in all species, but the positions and lengths of introns vary greatly, even as the number of introns remains mostly consistent. Correlations of gene structure among species pairs were low (fig. 6B); only 1% of introns are shared among all species (see the inner plot in fig. 6B), and 64.3% of the 167,873 introns from single-copy orthologs are unique. While several previous studies have demonstrated a high rate of intron loss in nematodes (Kent and Zahler 2000; Cho et al. 2004; Kiontke et al. 2004; Coghlan and Wolfe 2004), others focusing on highly conservative genes have reported much higher level of conservation in exon/intron structure (Ma et al. 2022). Our work, which includes a small set of species that have nearly complete genomes, reveals very low levels of gene structure conservation, and that even within an examination of a small fraction of known Clade V nematodes.

**Fig. 6.**
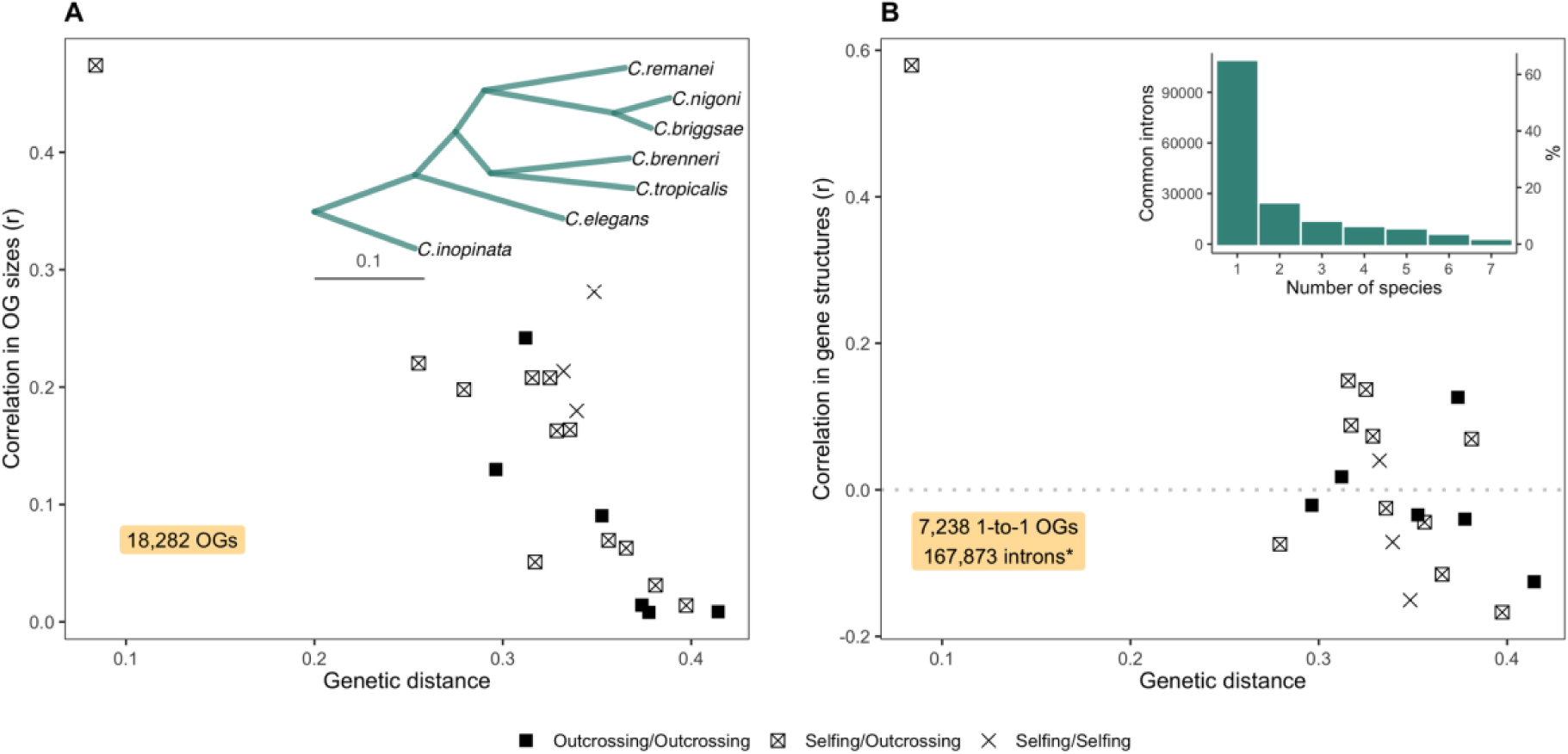
Correlations of orthogroup sizes and gene structures with genetic distances. **(A)** The correlation of orthogroup sizes across 18,282 orthogroups against the phylogenetic distance between species, using only orthogroups present in both species. The inset plot displays the phylogenetic tree from orthoFinder analysis, used for estimating genetic distances. Black boxes show comparisons of outcrossing species versus outcrossing species, boxes with crosses represent selfing versus outcrossing species, and crosses signify selfing versus selfing species. **(B)** The correlation of exon/intron gene structure across 7,238 single-copy orthologs present in all species. * We used 167,873 introns determined by GenePainter analysis, which includes only introns surrounded by coding sequences, without 5’ UTR introns. The inset plot shows a histogram illustrating the frequency of common introns across nematode species

We compared gene, exon, and intron length distributions, as well as the number of exons, among both all genes and single-copy orthologues genes in *Caenorhabditis* nematodes. Fig. 7A illustrates the mean and median values of exon and intron lengths, and supplementary fig. S7 (Supplementary material) the gene lengths for corresponding species, with the effect sizes (Cohen’s d) and significance from the non-paired Wilcoxon tests in 1-to-1 orthologues provided in supplementary fig. S8 (Supplementary material). Overall, these seven nematodes have very similar distributions in sizes of genes, exons, and introns and the number of exons (fig. 7B), despite substantial differences in effective population size across species. Further, all species have more introns in 1-to-1 orthologs than the average observed across all genes. *C. elegans* and *C. inopinata* tend to have longer genes and introns (supplementary fig. S8A and C, Supplementary material), and *C. nigoni* has fewer introns than other species (supplementary fig. S8B, Supplementary material). It is important to note that the quality of annotation in these species may impact these statistical results.

**Fig. 7.**
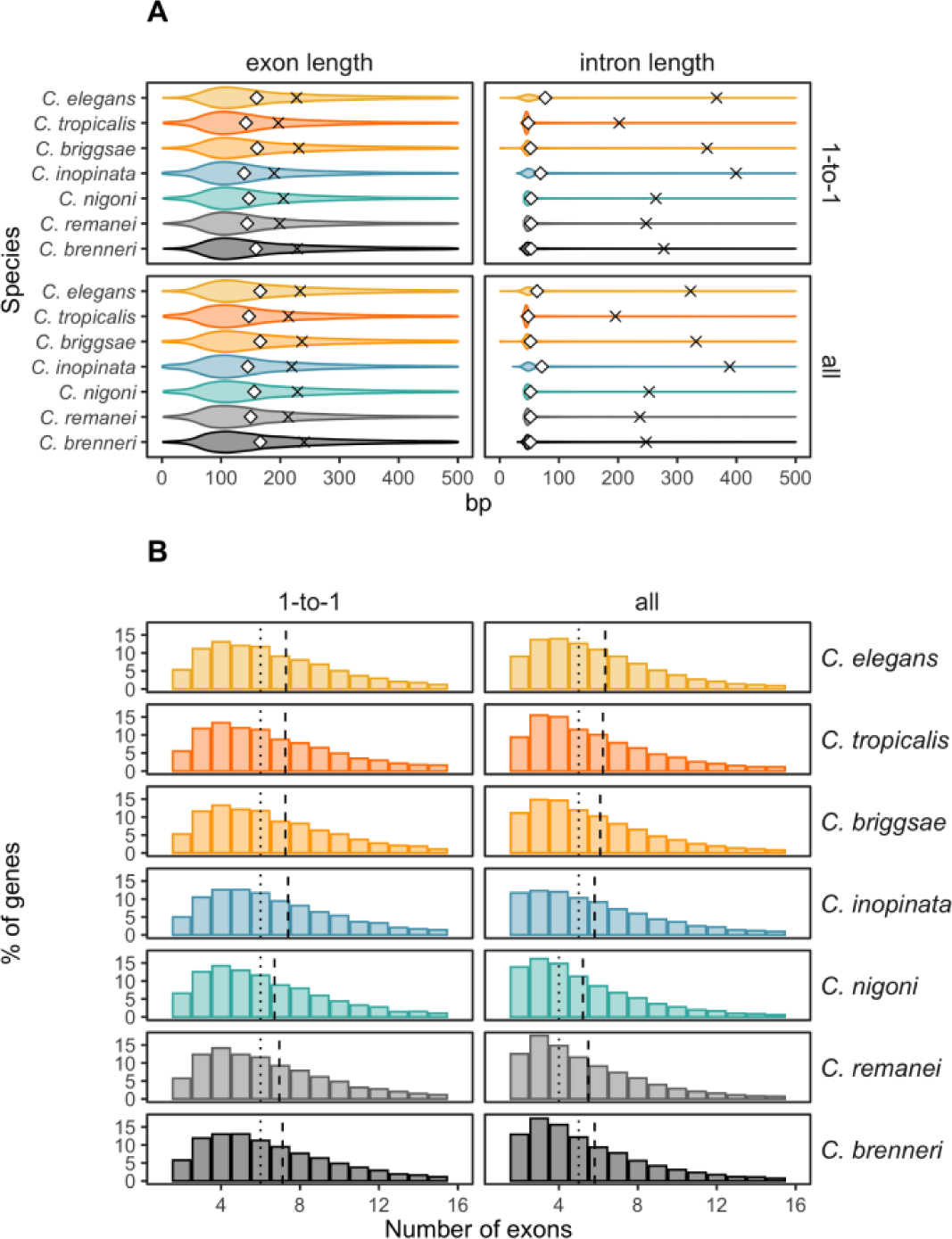
Distributions of genetic features in all genes and single-copy orthologs. **(A)** Exon and intron lengths in single-copy (1-to-1) orthologs and all genes within the orthogroups of the species. The crosses represent the mean values, and the diamonds represent the medians. **(B)** The number of exons in 1-to-1 orthologs and all genes within orthogroups. The dashed vertical lines indicate the means, and the dotted vertical lines indicate the medians.

Differences between selfing (*C. elegans*, *C. briggsae*, *C. tropicalis*) and outcrossing species (*C. brenneri*, *C. remanei*, *C. nigoni*, *C. inopinata*) in those gene features are summarized in Table 2. When comparing all genes, selfing species tend to be slightly longer genes and have approximately one more intron than outcrossing species. However, there are no discernible distinctions in gene, exon, and intron lengths, as well as exon numbers, when comparing these features in 1-to-1 orthologs between species with selfing and outcrossing reproductive modes and, thus, drastically distinct population sizes.

**Table 2.** Differences in gene features in selfing and outcrossing species. We used a linear mixed-effects model (lmer), with the reproduction system as a fixed effect and species as a random effect. The values in the columns represent p-values associated with the reproduction system, after the Bonferroni correction.

|  | Selfing vs Outcrossing species |  |  |  |  |  |
| --- | --- | --- | --- | --- | --- | --- |
|  | 1-to-1 orthologs |  |  | All genes |  |  |
| | mean ± standard deviation in selfers; outcrossers | Cohen's D | Linear model p-value, Bonferroni correction | mean ± standard deviation in selfers; outcrossers | Cohen's D | Linear model p-value, Bonferroni correction |
| Gene lengths | 3508.7 ± 4065.58; 3253.6 ± 3362.65 | 0.069 | 1 | 2891.8 ± 3763.76; 2523.2 ± 2914.29 | 0.113 | 1 |
| Exon lengths | 218.5 ± 235.79; 205.5 ± 225.23 | 0.057 | 1 | 227.9 ± 257.28; 225.9 ± 277.16 | 0.007 | 1 |
| Intron lengths | 306.4 ± 941.55; 299 ± 685.65 | 0.009 | 1 | 282.2 ± 914.22; 277.7 ± 653.87 | 0.006 | 1 |
| Exon number | 7.3 ± 4.73; 7 ± 4.51 | 0.049 | 1 | 6.2 ± 4.38; 5.6 ± 4.1 | 0.157 | 0.1303061 |

We used the alignments of exon/intron structures of 1-to-1 orthologs to compare certain features of shared introns, including length, phase, and splicing sites, across pairs of species, and between selfing and outcrossing nematodes. Interestingly, sister species *C. elegans* and *C. inopinata* on average longer have introns than other species (Supplementary fig. S9A, Supplementary material). Using a linear mixed-effects model, we do not find evidence that either phylogenetic distance nor reproductive mode have a substantial influence on the length differences of shared introns.

In addition to intron size, we compared the phases and splicing sites of introns shared across nematode species, as well as those unique within each species. Phase refers to how “perfect” the codon information is on either side of the intron (e.g., phase 0 means that the intron begins precisely at the end of the preceding codon). All species have more than 50% of introns in phase 0 (fig. 8A), consistent with the general observation that phase 0 introns are the most frequent in most organisms (Fedorov et al. 1992; Long et al. 1995; Nguyen et al. 2006; Rogozin et al. 2012), potentially due to their role in alternative splicing. *C. elegans*, *C. briggsae*, *C. inopinata*, and *C. nigoni* showed a significant difference (Fisher’s test) in the distribution of introns between phase 0 and non-phase 0 (combining phase 1 and phase 2) for the species-specific introns (“Unique” in fig. 8). In contrast, *C. brenneri* exhibited a significant difference between species-specific and common introns, with more non-phase 0 introns in those shared with other species. The majority of splicing sites of introns in 1-to-1 orthologs are the canonical GT-AG sites. We also compared variation in the fraction of other rare splicing sites between species-specific and shared introns among nematodes (fig. 8B), find no significant differences for the majority of species (Fisher’s test), except for the selfing *C. briggsae*, which exhibited more frequent rare types of splicing sites in species-specific introns (analysis of phase-based data can be found in Supplementary fig. S9B and C in Supplementary material).

**Fig. 8.**
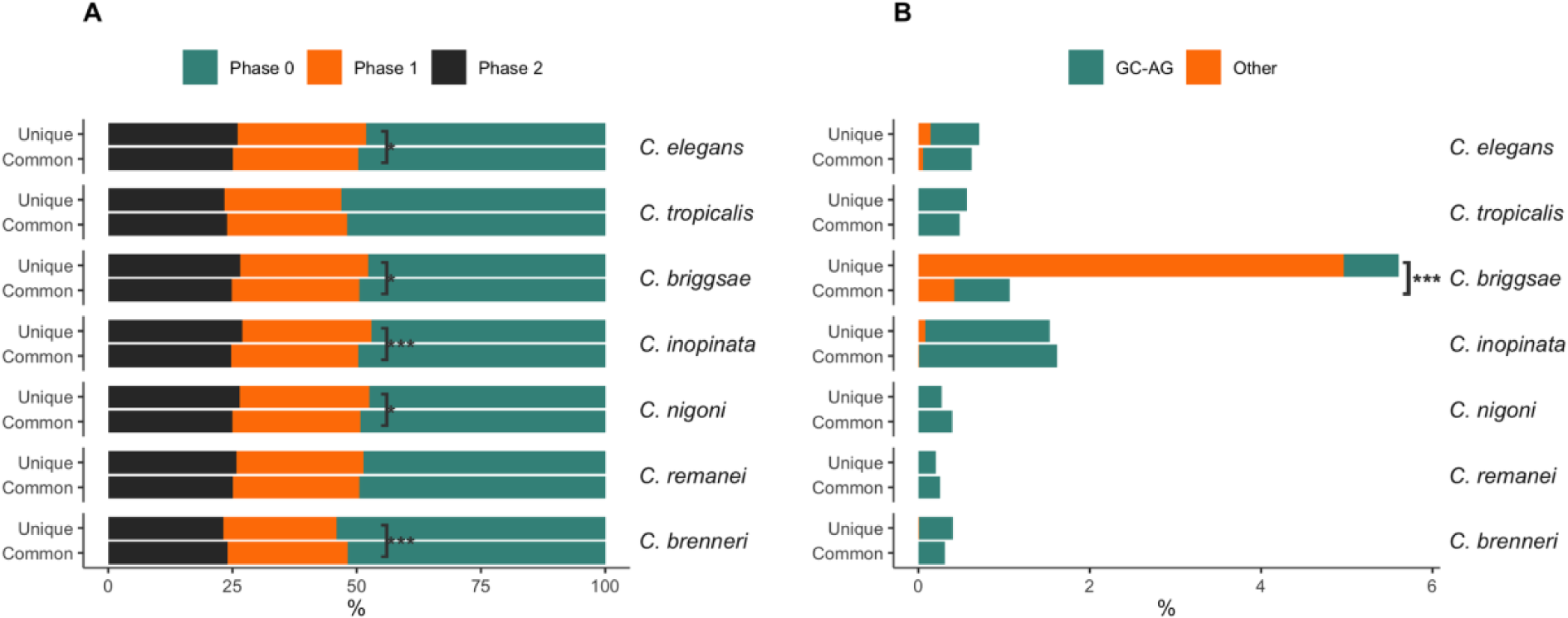
Intron features in species-specific (unique) introns and introns shared with other species (common). **(A)** Phases of introns. Colors represent the relative proportion of Phase 0 introns (teal), which means that the intron is located between codons, Phase 1 (orange), and Phase 2 (black). Asterisks indicate the statistical significance of differences in proportions of Phase 0 and non-Phase 0 (combined Phase 1 and Phase 2) introns between unique and common introns, tested using Fisher’s test after Bonferroni correction. * represents adjusted p-value <0.1; *** represents adjusted p-value <0.0001. **(B)** Relative percentage of splicing sites of introns. Only splicing sites other than the most common and canonical GT-AG are shown. GC-AG sites are depicted in teal, and all other rare types are shown in orange. The asterisk denotes the statistical significance of the difference in the fraction of sites other than GT-AG between unique and shared introns, with p-values from Fisher’s test with Bonferroni correction. *** represents that the adjusted p-value is <0.0001.

## Discussion

Nematodes provide a valuable metazoan model to study the impact of effective population size and reproductive system on the gene and genome organizations. *C. brenneri* is currently considered the most genetically diverse among all *Caenorhabditis* nematodes, and among the most diverse known metazoans (Dey et al. 2013), and completion of a high-quality genome and annotation for this species greatly expands the total scope of variation across this genus. The chromosome organization of *C. brenneri* follows the typical pattern observed in nematodes, featuring conservative macrosynteny across species, and large central domains with high gene density and low repeats density that exhibit more conservation than peripheral domains. Selfing species have lost more protein coding and noncoding DNA relative to outcrossing species, however all seven *Caenorhabditis* species have low correlation in orthogroup sizes and very high rates of gene turnover. Somewhat surprisingly given the general pattern of synteny seen at a macro scale, gene exon/intron structure at the individual gene level vary substantially between species, as they share just 1% of introns in single copy orthologs. Moreover, there is no significant differences in distributions of intron number or length in species with distinct population sizes or reproduction modes, suggesting underlying functional constraints in their evolutionary dynamics.

### Chromosome-level assembly of C. brenneri

The new complete genome of *C. brenneri* generated here exhibits much greater continuity, featuring complete chromosomes, and scant redundancy compared to the previously available genome of *C. brenneri* (generated by the Caenorhabditis brenneri Sequencing and Analysis Consortium), which was fragmented into thousands of scaffolds and highly duplicated due to the high residual heterozygosity within the sequenced strain. Given the extreme level of diversity observed within this species, the development of the highly inbred strain (CFB2252) was essential in achieving the desired quality and completeness of the assembled genome given the high level of diversity in this species. In addition to using long and short reads for assembly and Hi-C data for scaffolding, we also used high-accuracy long-read data from a single nematode. This approach suggests that single-individual long-read sequencing can be adapted to explore population-level variation in *C. brenneri* (and other species with substantial genetic diversity) to overcome challenges with mapping short-read data within populations that essentially have no single “reference” genome. The completeness of the *C. brenneri* genome was necessary for conducting comprehensive comparative analyses across nematode species, such as macro- and microsynteny, as well as genome and gene organization evaluations.

### Conservative macrosynteny and domain-like organization of the genome in C. brenneri

*C. brenneri,* like other rhabditid nematodes, displays notable stability of karyotype, five autosomes and one sex chromosome (Nigon 1949; Brenner 1974; De La Rosa et al. 2021). Syntenic reconstruction across the seven *Caenorhabditis* species with fully assembled genomes supports previous observations of multiple intrachromosomal inversions and rearrangements characterized by minimal interchromosomal changes but frequent intrachromosomal translocations and inversions (Stein et al. 2003; Hillier et al. 2007; Kanzaki et al. 2018; Teterina et al. 2020; Stevens et al. 2020; Sun et al. 2022). This conservative macrosynteny, characterized by fusion and intrachromosomal rearrangement of seven ancestral chromosome elements named Nigon elements, has been previously described throughout the nematode phylogeny (De La Rosa et al. 2021; Tandonnet et al. 2019). Currently, it has been proposed that the stability of nematode karyotype and such macrosynteny can be attributed to some combination of the control of chromosome pairing during meiosis and other mechanisms that regulate meiotic crossovers and genome integrity (McKim et al. 1993; Villeneuve 1994; Hillers and Villeneuve 2003; Lui and Colaiácovo 2013; Bhargava et al. 2020; Dokshin et al. 2020) and that potentially direct selection against interchromosomal translocation (Hillier et al. 2007).

Like other *Caenorhabditis* species, C*. brenneri* exhibits a domain-like organization for each chromosome with higher gene and lower repeat density in the central domains (The C. elegans Sequencing Consortium 1998; Parkinson et al. 2004; Cutter et al. 2009; Carlton et al. 2022; Woodruff and Teterina 2020; Stevens et al. 2020). Similarly, we also found greater conservation in the central regions of chromosomes, likely caused by similar patterns of suppression of recombination and linked selection observed in other *Caenorhabditis* species (Rockman and Kruglyak 2009; Ross et al. 2011; Teterina et al. 2023). Moreover, the central chromosomal domains have higher activity and more euchromatin compared to more inactive and heterochromatic arms in some *Caenorhabditis* species (Garrigues et al. 2015; Cabianca et al. 2019; Teterina et al. 2020; De La Cruz-Ruiz et al. 2023), potentially contributing to their greater level of conservation.

### Decay in microsynteny and lack of conservation in gene organization

Despite conservation of macrosynteny—most orthologous genes located on the same chromosome across species—nematodes display a striking lack of microsynteny (Coghlan and Wolfe 2002; Stein et al. 2003; Hillier et al. 2007; Kanzaki et al. 2018; Teterina et al. 2020; Stevens et al. 2020; Sun et al. 2022), with frequent rearrangements at the level of syntenic blocks and variable intron/exon structure at the level of individual genes. For example, we found that approximately half of the syntenic blocks are in reverse orientation, with the reversed blocks generally on average shorter than those in the forward orientation. Such structural changes in the genome are mirrored by a remarkably high rate of gene birth and death (Adams et al. 2023; Ma et al. 2024), resulting in low correlations in orthogroup sizes across the species studied here. Many gene families in nematodes are organized in clusters of co-located homologous genes, with some genes even being expressed together in an operon-like manner (Thomas 2006; Thomas and Robertson 2008; Chen and Stein 2006; Roy et al. 2002; Reinke and Cutter 2009; Cutter et al. 2009; Pettitt et al. 2014). Rapid gene turnover in nematodes is likely driven by an extreme rate of duplication (Woollard 2005; Lipinski et al. 2011; Farslow et al. 2015; Konrad et al. 2018), which makes it several times higher than the per-nucleotide substitution mutation rate (Denver et al. 2012; Denver et al. 2009; Saxena et al. 2019; Konrad et al. 2019). However, there is no clear relationship between rates of gene turnover and population size across species. Given the high rate of divergence in gene composition among species, analysis of variation within species is likely to be helpful to deepen our understanding of these processes.

At the level of individuals genes, nematodes are known to lose introns at a high rate (Kent and Zahler 2000; Stein et al. 2003; Logsdon 2004; Coghlan and Wolfe 2004; Cho et al. 2004; Kiontke et al. 2004; Roy and Gilbert 2005; Roy and Penny 2006; Ma et al. 2022). In particular, they tend to mostly lose short introns, usually precisely, more frequently on the 3’-end of the gene, and preferentially in genes with high expression (Ma et al. 2022). Indeed, as has been previously observed for nematode genomes, we find very low conservation in exon/intron structures of 1-to-1 orthologues, with only 1% of common introns in seven species with complete genomes compared here (Coghlan and Wolfe 2004; Cho et al. 2004); in contract, mammals exhibit nearly-complete conservation of intron positions (Roy et al. 2003; Coulombe-Huntington and Majewski 2007; Poverennaya et al. 2020). An apparent exception to this pattern is the somewhat conserved pattern of intro structure for extremely conserved single-copy orthologs that show very slow rates of evolution across nematodes as a whole (Ma et al. 2022). One mechanism that could explain the dynamic nature of exon/intron evolution could potentially be the high level of gene birth and death in nematodes, which would allow new introns to arise and disappear in paralogs in a somewhat cryptic fashion. This pattern has been described for a subset of gene families within these species (Robertson 1998; Cho et al. 2004). The answer to the balance between macro conservation and micro genomic churn is likely to be better resolved as we build more high-quality assemblies at a variety of phylogenetic scales that allow microsyntentic relationships to be assessed with adequate confidence.

Overall, our data reveals an apparent conundrum: introns apparently turn over in location in a very dynamic fashion within genes, yet our major observation is that the distribution of intron number and length across the entire genome is very similar for each species. How can both be true? The answer may lie in the functional role that introns play: mRNA processing and expression. Specifically, splicing of introns plays a critical role in facilitating the export of mRNA from the nucleus to the cytoplasm (Shiimori et al. 2013; Katahira 2015; Zheleva et al. 2019; Xie and Ren 2019). Most importantly for this discussion, introns regulate gene expression in nematodes, especially when they are located on 5’-end of the gene (Bieberstein et al. 2012; Crane et al. 2019; Sands et al. 2021), with the insertion of even a single intron increasing gene expression by as much as 50% (Crane et al. 2019). Moreover, trial and error in *C. elegans* molecular biology has shown that it is best to use at least three introns when creating artificial transgenes in order to achieve an optimal level of expression (Fire 1995). Interestingly, this is close to the average number of introns in all species (fig. 7B). Furthermore, introns can regulate expression of genes through the alternation of chromatin marks (Bieberstein et al. 2012; Jo and Choi 2019), as well as generating biased allele-specific expression (Sands et al. 2021).

Efficiency of intron splicing depends on the nature of the splicing sites themselves, and nematodes have U2 type of spliceosomes that only require a short motif for site recognition (Wahl et al. 2009; Bartschat and Samuelsson 2010). Roughly half of introns across all studied species were in Phase 0, such that they do not disrupt a codon. This is likely driven by a need to preserve codon integrity during alternative splicing, in keeping with studies in other non-nematode species (Fedorov et al. 1992; Long et al. 1995; Nguyen et al. 2006; Rogozin et al. 2012). Overall, the global hypothesis most consistent with the pattern of intron structure across the broad diversity of these species is that introns themselves are functionally important and clearly under selection (no real signal of population size on intron evolution), but it is the presence of introns that is important and not necessarily their precise placement within a gene.

### Genome organization in selfing and outcrossing species

An early observation regarding genomic evolution in *Caenorhabditis* was that the genomes of selfing nematodes tend to be substantially smaller than genomes of their outcrossing relatives. This 10-30% reduction is size turns out to be mostly attributable to extensive loss of genes and flanking regions (Bird et al. 2005; Fierst et al. 2015; Yin et al. 2018; Stevens et al. 2019; Adams et al. 2023). Reduction of some regions of the genome can in fact can be driven by adaptation to self-fertilization itself rather than an epiphenomenon of the reduction in effective population size under selfing. For instance, in *C. briggsae* and other selfing species some genes with male-biased expression such as sperm competition proteins seem to have been preferentially lost Yin et al. 2018). Furthermore, hybrids of *C. briggsae* and *C. nigoni* show sex-biased gene misregulation, especially in spermatogenesis (Sánchez-Ramírez et al. 2021). However, another explanation for the difference in genome sizes between selfing and outcrossing species that has been suggested is the expansion and duplication coding genes in the genomes of outcrossing species (Kanzaki et al. 2018; Stevens et al. 2019; Adams et al. 2023). However, we found only minor changes in the size of shared orthogroups, with outcrossing species tending to have more species-specific orthogroups.

A final explanation for the differences in genome sizes arises from an interaction of genome-level mutational processes and population genetics. In particular, it has been demonstrated that deletions are more common than insertions in *C. elegans* and that these deletions are predominantly located in intergenic regions as well as being more frequently found on chromosome arms (Hwang and Wang 2021; Zhang et al. 2022). Reduced population size and a very low effective recombination rate within selfing species means that deletions have a higher probability of fixing than in outcrossing species (Thomas et al. 2012; Fierst et al. 2015; Hartfield et al. 2017; Hartfield 2016). With our new genomic data, we also find that the main distinction between selfing and outcrossing species is in the fraction of intergenic DNA and number of exons/genes (Fierst et al. 2015; Stevens et al. 2019; Adams et al. 2023). While all of these processes are likely to be involved in driving genomic evolution across these species, we need to move toward a full population genomics of independently assembled genomes segregating within natural populations to tease them apart any more fully than the current state of descriptive comparison among species.

### Testing the hypothesis that effective population size structures the evolution of genomic organization

The species included here range from those with little to no polymorphism within populations (*C. tropicalis*) to polymorphism at the level of one SNP every 10 base pairs (*C. brenneri*). Surely if population size is the main driver of the evolution of many genomic features, then we should see that signature here (Lynch 2002; Lynch and Richardson 2002; Lynch and Conery 2003; Kiontke et al. 2004). Despite this expectation, we find that the distributions of gene, exon, intron lengths, and exon number are mostly similar across all species, and no significant differences are detected found either between selfing and outcrossing species or among outcrossing species with orders of magnitude differences in effective population sizes. Taken together, the observation of frequent intron loss and the stability of the number of introns across species with very different population sizes suggests instead that natural selection on the balance between loss and functional necessity of introns best explains these patterns. And while the influence of population size on the evolution of putatively “nearly neutral” genetic elements has certainly been an important feature of many theories of molecular evolution, it has also been simultaneously suggested that once these features come to be present within genomes, they will quickly evolve other functional roles (Lynch 2007; Jo and Choi 2015; Girardini et al. 2023) The molecular biology of *Caenorhabditis* nematodes, including truly hyper-polymorphic species, suggests that the latter circumstance dominates the former and that most features of the genome are under direct selection, even if it is not selection on a specific sequence of DNA.

## Materials and Methods

### Strain generation and sequencing

The *C. brenneri* line CFB2252 was initially isolated as a single gravid female from a rotting marigold flower in Shyamnagar, India. The line was propagated though sib-mating under standard nematode lab-rearing conditions (Brenner 1974) for twenty generations with population expansion every five generations to help purge the accumulation of large-effect deleterious mutations. For the next twenty-two generations a single gravid female was picked for propagation of the strain. The population was expanded again for eight generations by chunking multiple worms each next generation. Finally, a line was generated by single-female propagation for another forty generations before being frozen. Thus, the final sequencing line was inbred for a total of ninety generations.

Genomic DNA and total RNA were isolated from a mixed stage population using the Zymo Genomic DNA Clean and Concentrator-25 (DCC) kit following Proteinase K digestion and the Zymo DirectZol RNA microprep kit (R2062) following three freeze/thaw steps in trizol. Genomic DNA from individual day one virgin female worms was used directly from Proteinase K digested samples. For long read sequencing we utilized the PacBio CLR sequencing with Sequel I platform with genomic DNA from a mixed stage population and for single worm PacBio we utilized the HiFi genomic (CCS) platform. For the PacBIo IsoSeq (transcript isoform) sequencing, we submitted total RNA. For short read sequencing we utilized the Illumina HiSeq 4000 platform for both genomic DNA and RNA sequencing prepared as mentioned above. 100ng of total RNA was used for mRNA extraction and library preparation with the Kapa mRNA Hyper prep kit (KK8580). Similarly, Hi-C sequencing using the Phase Genomics platform utilized genomic DNA prepared as above. All sequencing was performed at the University of Oregon Genomics and Cell Characterization Core facility (GC3F) (https://gc3f.uoregon.edu/).

### Genome assembly

Potential adapters sequences were removed from each long read using removesmartbell.sh from bbmap v.38.16. Hi-C read and other short reads were trimmed and filtered using skewer v.0.2.2 (Bushnell 2014; Jiang et al. 2014). Genome assembly was conducted with PacBio CLR reads by FALCON v.1.2.3 and FALCON-Unzip v.1.1.3 (Chin et al. 2016), followed by additional phasing with Hi-C reads by FALCON-Phase v.1.0.0 (Kronenberg et al. 2021). Contigs were polished with Illumina short reads by Pilon v.1.23 (Walker et al. 2014) with bwa v.0.7.17 (Li 2013), and samtools v.1.5 (Li et al. 2009). Then we performed contig scaffolding with Hi-C data following the 3D-DNA pipeline (Dudchenko et al. 2017) with run-asm-pipeline.sh v.180922 and Juicer v.1.6.2 (Durand et al. 2016), bwa and samtools. Then scaffolds were polished with HiFi reads by RACON v1.4.20 (Vaser et al. 2017). HiFi reads were corrected using Illumina read with fmlrc v1.0.0 (Wang et al. 2018). In addition, we assembled HiFi reads from an individual nematode of the reference strain (CFB2252) using hifiasm v.0.15.1-r334(Cheng et al. 2021), and assembled short read obtained from the pool of individual by SGA v.0.10.15 (Simpson and Durbin 2012). Contigs from those two assemblies were used for validation of the final genome assembly. Gaps in the *C. brenneri* genome were filled using corrected and uncorrected HiFi reads by LR_GapCloser (Xu et al. 2019) and TGS_GapCloser (Xu et al. 2020). We assembled and annotated a mitochondrial genome from HiFi reads with MitoHiFi v3.0.0 (Uliano-Silva et al. 2023) using the mitochondrial genome of *C. brenneri* as a reference (the accession number in Gene Bank is NC_035244.1). We removed bacterial contamination using bloobtools v1.1.1 (Laetsch and Blaxter 2017), and validated scaffolds using HiFi reads with Inspector v1.0.1 (Chen et al. 2021). The quality of the final genome assembly was accessed with Inspector, MERQURY v2020-01-29 (Rhie et al. 2020), BUSCO v5.4.2 (Simão et al. 2015) with databases nematode_odb10 and metazoa_odb10, and coverage from HiFi, Illumina reads, hifiasm and sga assemblies using pbmm2 from pb-assembly v1 (https://github.com/PacificBiosciences/pb-assembly), samtools, and bedtools v2.25.0 (Quinlan and Hall 2010). The heterozygosity was estimated for the HiFi long reads from the individual nematode from the CFB2252 strain with jellyfish v2.2.10 (Marçais and Kingsford 2011) and GenomeScope v2 (Vurture et al. 2017).

### Genome annotation and repeat masking

We masked repetitive sequences in *C. brenneri* genome with a approach described in detail in Teterina et al. (2020), with RepeatMasker v.4.0.7 (https://github.com/rmhubley/RepeatMasker), RepeatModeler v.1.0.11 (http://www.repeatmasker.org/RepeatModeler/), detectMITE v.2017-04-25 (Ye et al. 2016), transposonPSI (http://transposonpsi.sourceforge.net), LTRharvest and LTRdigest from GenomeTools v.1.5.11 (Gremme et al. 2013), tRNAscan v.1.3.1 (Lowe and Eddy 1997), Blast .2.2.29+ (Altschul et al. 1990), and USEARCH v.8.0 (Edgar 2010). We annotated the genome using RNA-seq data and single-molecule long-read RNA sequencing (Iso-Seq), and protein sequences of *C. briggsae* (PRJNA10731), *C. brenneri* (PRJNA20035), *C. elegans* (PRJNA13758), and *C. remanei* (PRJNA577507) from WormBase 287 and *C. wallacei* (strain JU1898) and *C. doughertyi* (strain JU177) from the Caenorhabditis Genomes Project v.2 (http://download.caenorhabditis.org/v2/).

We trimmed RNA-seq reads by skewer and mapped them to the *C. brenneri* genome with STAR v2.5.3 (Dobin et al. 2013), these alignments were used as evidence for annotation with BRAKER v2.1.0 (Hoff et al. 2019). Then, we used *Caenorhabditis* proteins and BRAKER gene predictions to perform annotation with MAKER. v.2.31.9 (Cantarel et al. 2008). The Iso-Seq data was processed using the isoseq3 pipeline (https://github.com/ylipacbio/IsoSeq3). Then annotation from Braker, Maker and collapsed isoforms from isoseq3 pipeline were combined by Evidence Modeler v.1.1.1 (Haas et al. 2008). Then we leveraged splicing prediction using RNA-seq alignments with the PASA pipeline v2.5.3 (Haas 2003). We validated the final assembly with RNA-seq reads, blast to nematode proteins, and functional annotation with Interproscan v. 5.27-66.0 (Jones et al. 2014), and BUSCO. Several genes were filtered as their exons were located in the masked regions of the genome and classified as repetitive elements by RepeatClassifier from RepeatModeller. We updated gene features and formatted annotation files using AGAT tools v1.0.0 (https://github.com/NBISweden/AGAT) and custom scrips provided in the GitHub repository for this project.

### Comparative genomic analysis

#### Divergence and conservation

We aligned the new soft-masked genome of *C. brenneri* with genomes of *C. elegans* (PRJNA13758), *C. remanei* (PRJNA577507), and *C. tropicalis* (PRJNA53597) from WormBase WS287, and *C. wallacei*, *C. doughertyi*, *C. sp44*, *C. sp48*, *C. sp51*, and *C. sp54* from the Caenorhabditis Genomes Project v.2 using progressiveCactus (Armstrong et al. 2020). The alignments were splitted to 50-kb windows by hal2maf_split.pl from HAL tools (Hickey et al. 2013), converted to MSA format with msaconverter (https://github.com/linzhi2013/msaconverter) and filtered by maffilter (Dutheil et al. 2014). The maximum-likelihood trees were constructed using IQ-TREE v2.2.0.3 (Nguyen et al. 2015). We got topologies of the trees, a total lengths of *C. brenneri* and *C. sp48* clade or solely the *C. brenneri* branch from a few trees where they were not clustered together. The total tree heights were estimated using package ape v5.3 (Paradis et al. 2004) in R v3.5.0 (R Core Team 2018).

We conducted a conservation sequence analysis using progressiveCactus alignments, which were converted from HAL to MAF format using hal2maf from the HAL tools, and utilized phyloFit and phastCons tools from Phast v1.5 (Siepel et al. 2005). First, we generated an initial model with phyloFit using the following tree “(((((*C. brenneri*, *C. sp48*), (*C. sp44*, *C. sp51*)), (*C. doughertyi*, (*C. tropicalis*, *C. wallacei*))), *C. remanei*), *C. elegans*, *C. sp54*)” and the whole genome alignments. Following that, we retrained the model for each chromosome with phastCons, and finally, applied these models to estimate conservation in 50-kb window alignments for the corresponding chromosomes and calculated the average scores within those windows. The visualization was done in R with packages ape, ggplot2 v3.4.2 (Wickham 2016), ggtree v1.14.6 (Yu et al. 2017), and patchwork v1.1.1 (Pedersen 2024).

#### Synteny, orthology, and gene structures

For orthology analysis, we used six *Caenorhabditis* species with chromosome-scale genome assemblies: genomes and canonical annotations of *C. elegans* (PRJNA13758), *C. remanei* (PRJNA577507), *C. inopinata* (PRJDB5687), *C. briggsae* (PRJNA10731), *C. nigoni* (PRJNA384657) were obtained from the WormBase WBPS18 and the genome and annotation for NIC58 strain of *C. tropicalis* were sourced from (Noble et al. 2021). The corresponding protein sequences were generated from the longest transcripts by AGAT tools. Orthology analysis was conducted by Orthofinder v2.5.5 (Emms and Kelly 2019). Then we estimated the number of exons, lengths of genes, exons, and introns without all genes that have orthogroups and genes that are in single copy in all 7 species (1-to-1) orthogroups. We used R packages stats v3.5.0, dplyr v1.0.7 (Wickham et al. 2022), tidyr v1.1.3 (Wickham 2021), esvis v0.3.1 (Anderson 2020), data.table v1.12.8 (Dowle et al. 2021), reshape2 v1.4.4 (Wickham 2007), ggplot2, corrplot v0.90 (Wei and Simko 2021), RColorBrewer v1.1.2 (Neuwirth 2014), ggplotify v0.0.8 (Yu 2021), gridGraphics v0.5.1 (Murrell and Wen 2020), patchcwork to compare and visualize these distributions and difference between species. The specific tests are indicated in figure legends and the main text.

The inference of syntenic blocks was performed based on the results from Orthofinder2 by R package GENESPACE v1.2.3 (Lovell et al. 2022). We visualized the riparian plot with *C. brenneri* as the reference genome, and estimated the number of genes, exons, introns, and fractions of exons/introns within corresponding syntenic blocks. As some species were lacking the annotation of the 3’ and 5’ UTR and other, we called regions that were neither exons nor introns as ‘intergenic.’ We then calculated the ratios of these measurements for species pairs in relation to *C. brenneri*. This allowed us to explore the connection between genome organization and the differences in population size ratios among species, which sometimes span orders of magnitude. However, due to a lack of population size estimates for certain species like *C. nigoni*, we focused on the classification of selfers versus outcrossers using ratios of the number of exons, introns, and the length of exons, introns, and intergenic regions. We replicated syntenic blocks proportionally to their sizes in the *C. brenneri* genome and trained a generalized linear model (glm) and a gradient boosting model to determine whether the ratios correspond to selfing species (*C. elegans* and *C. tropicalis*) vs outcrossing *C. brenneri* or outcrossing species (*C. remanei* and *C. inopinata*) vs *C. brenneri*. Then we evaluated the model with selfing *C. briggsae* vs *C. brenneri* and outcrossing *C. nigoni* vs *C. brenneri*. For the generalized linear model (glm) and gradient boosting model (XGboost) approaches, we utilized R packages glmnet v2.0.18 (Friedman et al. 2010), xgboost v1.6.0.1 (Chen et al. 2022), tidyr, dplyr, caret v6.0.8 (Kuhn 2020), broom v0.7.9 (Robinson et al. 2021), pROC v1.18.0 (Robin et al. 2011), caTools v1.17.1.2 (Tuszynski 2019), gtable v0.3.0 (Wickham and Pedersen 2019), DALEX v2.4.3 (Biecek 2018).

#### Comparison of exon/intron gene structures

We explored the exon/intron structures of single copy orthologs determined by OrthoFinder2 and corresponding gene annotations using GenePainter v2.0 (Hammesfahr et al. 2013; Mühlhausen et al. 2015). Then we estimated the sizes of the introns, accessed phases of the CDS (Phase 0,1, or 2) that follows the intron, and obtained splicing sites using *agat_sp_extract_sequences.pl* form AGAT tools v1.2.0 to get splicing sites. Then we parsed sizes, phases, and splicing sites of introns with gene exon/intron structures alignments from GenePainter and compared unique and common introns as well as estimated association and difference of introns shared in pairs of nematode species using R packages dplyr, reshape2, lme4 v1.1.27.1 (Bates et al. 2015), lmerTest v3.1.3 (Kuznetsova et al. 2017), MASS v.7.3.54 (Venables and Ripley 2002), ggplot2, patchwork, ggplotify, gridGraphics, corrplot.

## Supplementary Materials

Supplementary figures (S1-S9) and Supplementary table S1 are available at *Molecular Biology and Evolution* online (Supplementary material).

## Acknowledgments

We thank Mark Blaxter and the Caenorhabditis Genomes team for permission to use the unpublished genome sequences for a number of species for some of our analyses. Asher Cutter kindly provided the semi-inbred progenitor of the CFB2252 line. The project was supported by funding from the National Institutes of Health to PCP (R35GM131838). We thank members of the Phillips laboratory and Kern-Ralph co-lab at the University of Oregon for the helpful discussions. The thank the University of Oregon Genomics and Cell Characterization Core Facility (GC3F) for support, and the Research Advanced Computing Services team at the University of Oregon for assistance with the high-performance computer Talapas.

## Data availability

Genomic and transcriptomic data generated in this work available at the European Nucleotide Archive database (ENA, https://www.ebi.ac.uk/ena) under BioProject ID PRJEB72296, the *C. brenneri’s* genome accession number is GCA_964036135.1. Copies of the current version of the genome and annotation of *C. brenneri* used in the analysis are available at Figshare under doi:10.6084/m9.figshare.25988629, all files necessary to estimate statistics and generate the figures available at Figshare under doi:10.6084/m9.figshare.25988626. Scripts for the analysis and visualization available at https://github.com/phillips-lab/Cbren_genome.

