## Supplementary Material for "Pervasive conservation of intron number and other genetic elements revealed by a chromosome-level genomic assembly of the hyper-polymorphic nematode *Caenorhabditis brenneri*"

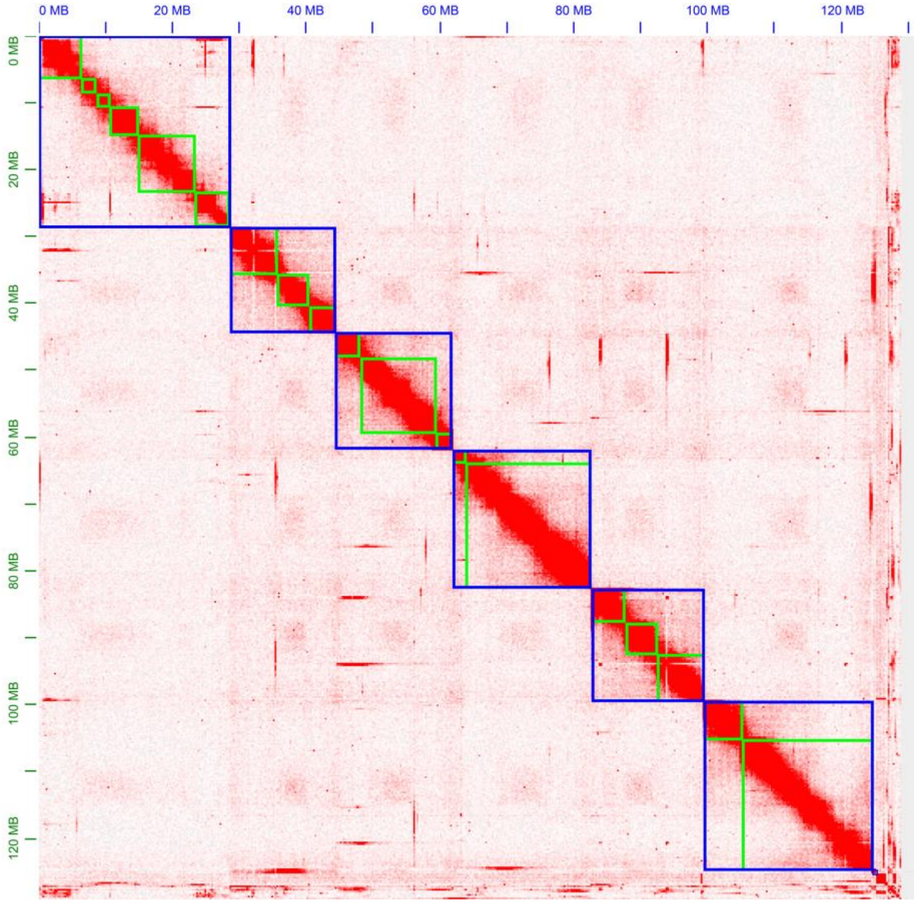


**Supplementary fig. S1.** Hi-C contact map generated with the 3D-DNA scaffolding pipeline and visualized with the Juicer tool. The x and y axes represent genome coordinates, with red color intensity indicating interaction frequency. Contigs are shown in green boxes, while blue boxes represent scaffolds.

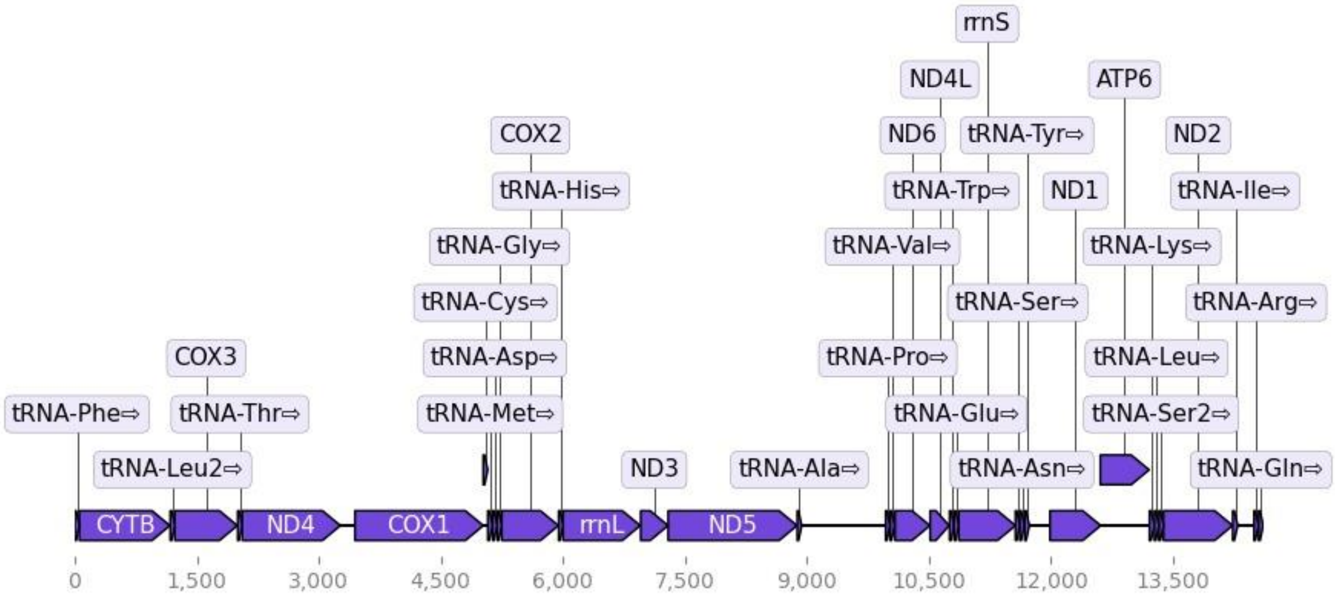


**Supplementary fig. S2.** Mitochondrial genome of *C. brenneri*. Predicted genes are labeled throughout the mitochondrial genome, maintaining the same order as the other available genome (NC_035244.1). The figure was produced by the MitoHiFi.

**Table S1.** Comparative synteny analysis with *C. brenneri*.

The syntenic blocks were determined be the GENESPACE analysis (see Methods). Two blocks (one in *C. briggsae* and one in *C. nigoni*) were removed from the analysis as they were small and located not in the same chromosome relative to *C. brenneri*.

| Focal species | Phylogenetic distance to *C. brenneri* (OrthoFinder) | Number of syntenic blocks with *C. brenneri* | Number of blocks in the same (reversed) orientation and [% in the same] | Longest syntenic block in *C. brenneri* / focal species, Mb | Mean size of the block in *C. brenneri* /focal species ±standard deviation, 100Kb | Mean size of the block in the same (reversed) orientation in *C. brenneri* / focal species, 100Kb |
|---|---|---|---|---|---|---|
| *C. tropicalis* | 0.255 | 102 | 62 (40) – [60%] | 23.3/12.8 | 11.2±25.9/7.51±15.4 | 14.1±32.2(6.76±9.56)/ 9.56±19.1 (4.33±5.07) |
| *C. remanei* | 0.312 | 131 | 61 (70) – [46%] | 14.6/17.1 | 8.34±15.6/8.72±16.9 | 10.6±21.2 (6.37±7.68)/ 11.0±23.0 (6.76±8.34) |
| *C. elegans* | 0.329 | 221 | 116 (105) – [52%] | 5.9 /3.9 | 5.00±7.96/4.23±5.64 | 5.23±7.54 (4.75±8.42)/ 4.58±5.87 (3.83±5.37) |
| *C. briggsae* | 0.336 | 130 | 65 (65) – [50%] | 11.6/9.1 | 8.65±16.0/7.89±11.8 | 12.2±21.7 (5.09±4.48)/ 10.4±15.6 (5.34±4.82) |
| *C. nigoni* | 0.353 | 132 | 70 (62) – [53%] | 14.6/12.8 | 8.45±17.0/8.59±15.4 | 11.1±22.4 (5.50±5.65)/ 10.8±20.2(6.05±5.88) |
| *C. inopinata* | 0.378 | 179 | 92 (87) – [51%] | 4.2/0.9 | 5.85±7.27/6.37±6.56 | 4.63±4.68(7.13±9.11)/ 5.83±6.18 (6.95±6.94) |

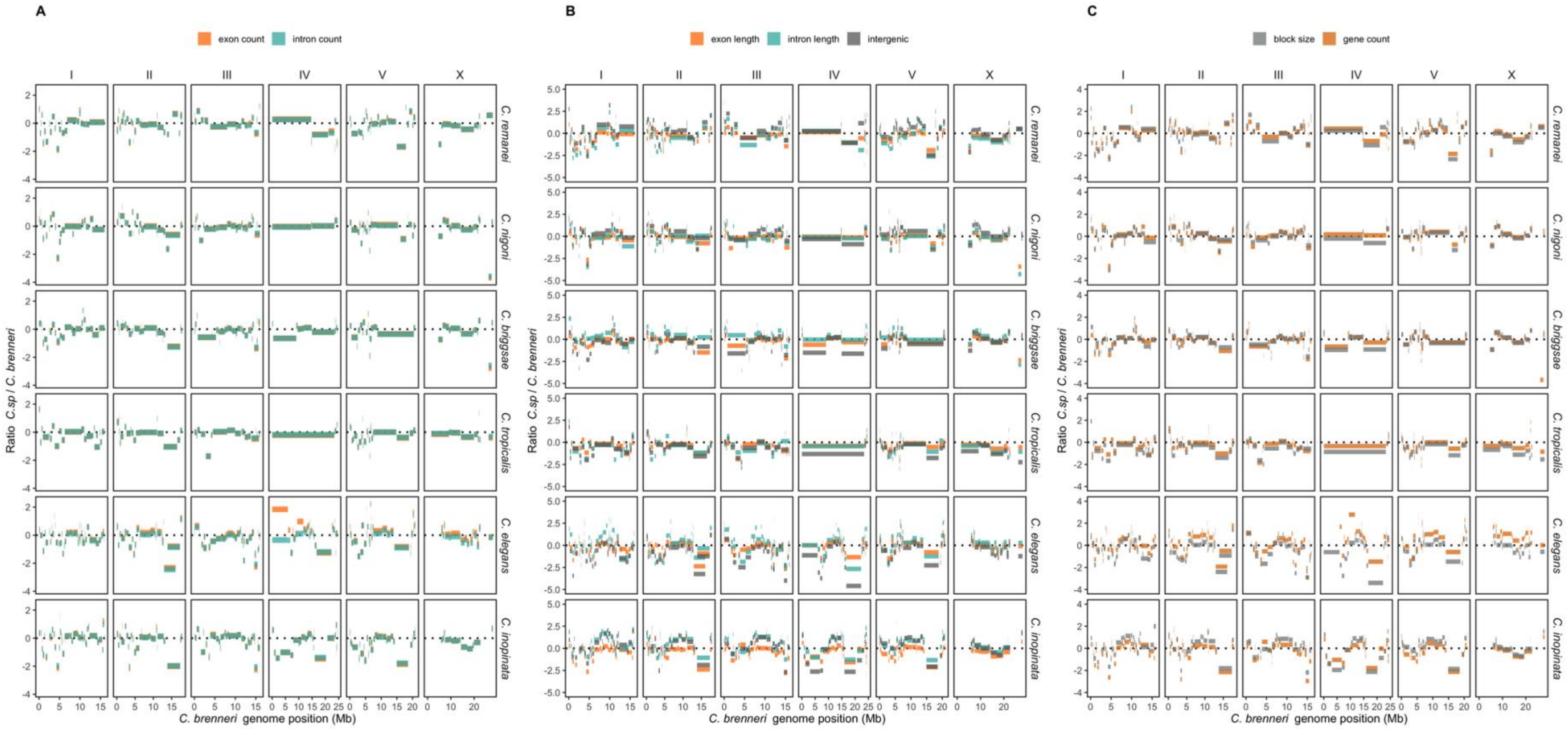


**Supplementary fig. S3** Ratios of genomic features of nematode species relative to *C. brenneri* in the corresponding syntenic blocks along the *C. brenneri* genome. **(A)** Ratios of exon and intron counts in focal species relative to *C. brenneri*. **(B)** Ratios of fractions of exons, introns, and intergenic regions that occupy syntenic blocks. **(C)** Ratios of syntenic block sizes and gene counts on them.

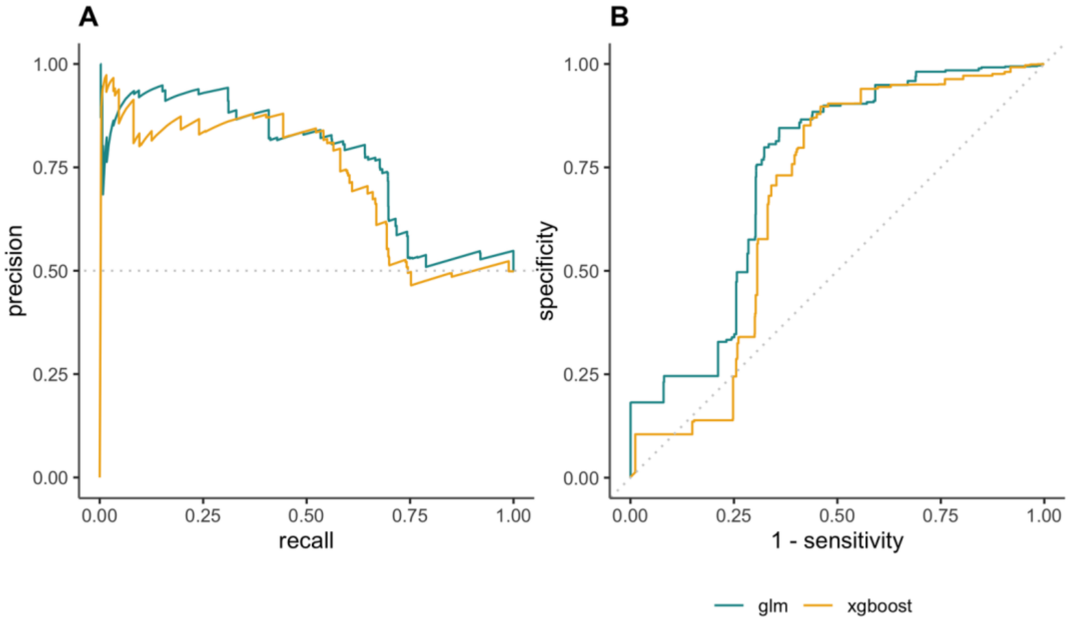


C

| | glm | xgboost |
|---|---|---|
| *AUROC* | 0.7363 | 0.6773 |
| *Accuracy* | 0.6792 | 0.6827 |
| *MAE* | 0.3545 | 0.3867 |
| *MSE* | 0.2323 | 0.2701 |
| *F1* | 0.505 | 0.5119 |

**Supplementary fig. S4**. Evaluation of classifications of selfing and outcrossing nematodes using glm and gradient boosting (xgboost) methods. **(A)** The precision-recall curve, illustrating how the model balances precision and recall across various threshold values. **(B)** The specificity to 1-sensitivity curve, demonstrating the model's capacity to identify true negatives. **(C)** A table with accuracy metrics: "AUROC" (Area Under the Receiver Operating Characteristic curve) shows the model's ability to differentiate between classes, "Accuracy" indicates the proportion of correctly classified instances, "MAE" (Mean Absolute Error) and "MSE" (Mean Squared Error) measure the average magnitude of errors, and the "F1" score represents the harmonic mean of precision and recall.

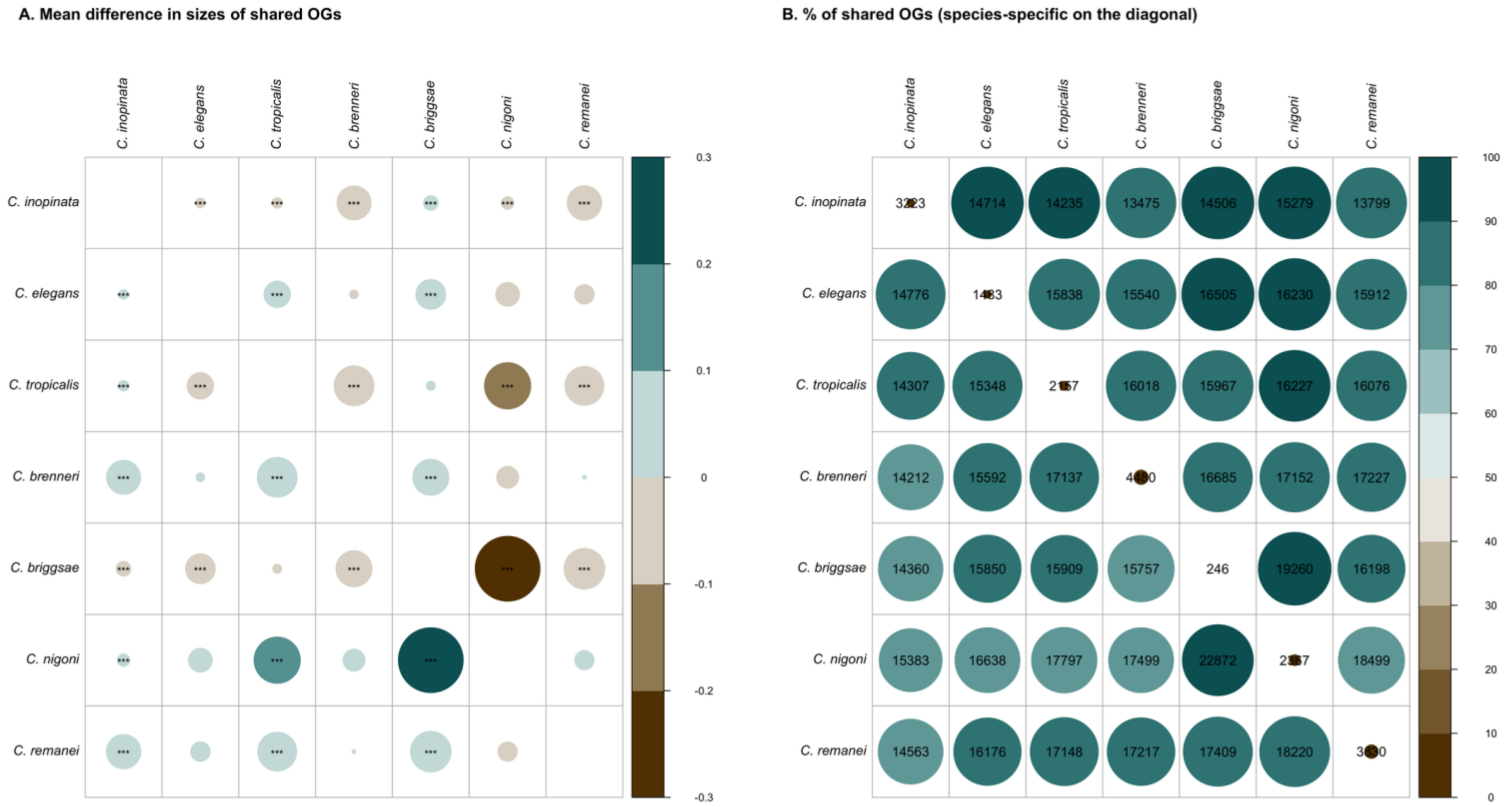


**Supplementary fig. S5**. **(A)** Mean differences in shared orthogroup (OG) sizes between species, and **(B)** the percentage of shared orthogroups between species, depicted by the color and size of circles. The numbers in each cell represent the number of genes a species has in orthogroups shared with another species. For instance, the cell in the second row and first column shows how many genes has *C. elegans* in orthogroups that are shared with *C. inopinata*. The diagonal showcases the number of species-specific genes.

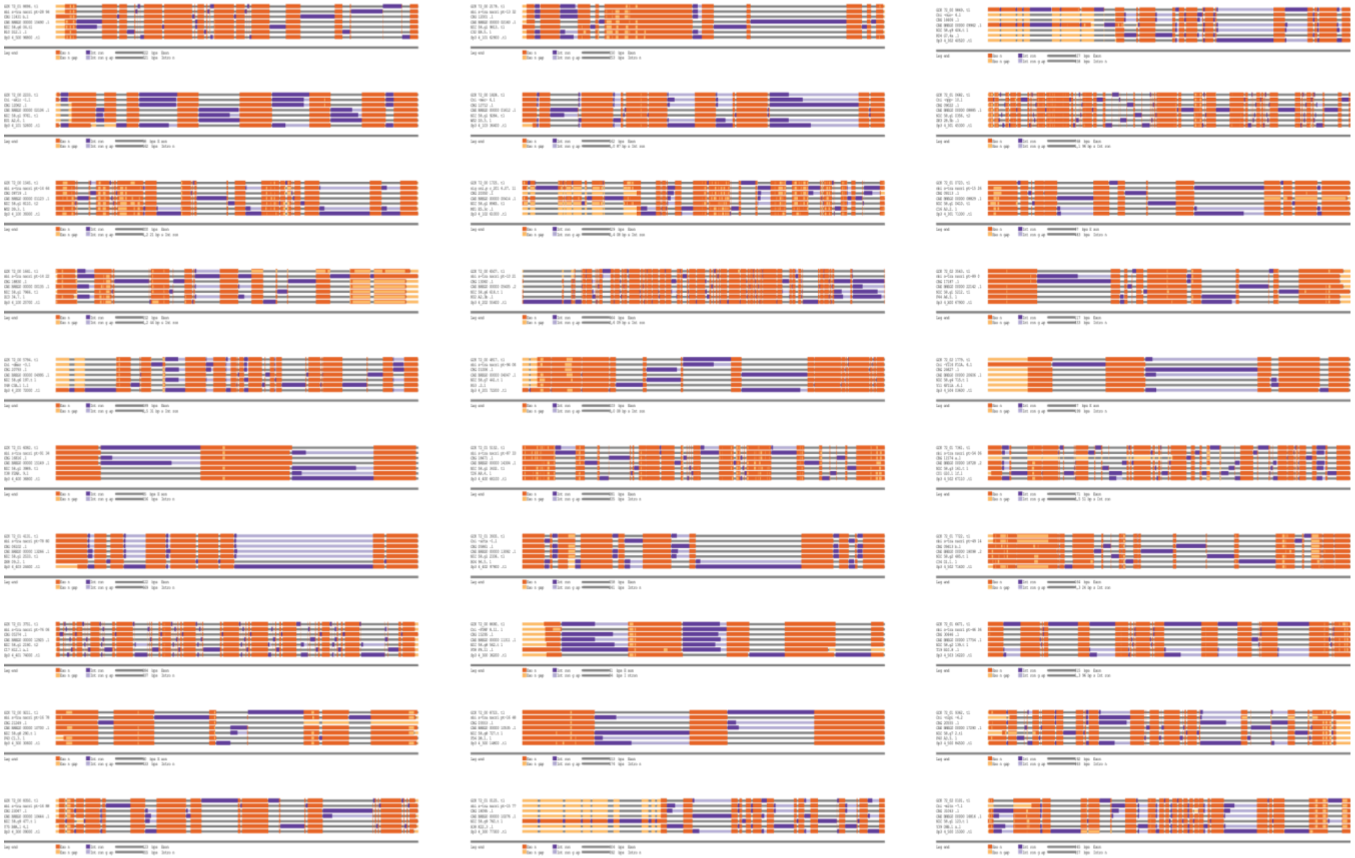

**Supplementary fig. S6**. Examples of exon/intron gene structure alignment for single-copy orthologs generated by GenePainter. Thirty randomly selected alignments, all scaled to the same length. The species order matches the one on fig. 4. Aligned exons are shown in orange, and light orange marks gaps in exon alignment. Aligned introns are shown in dark purple, with light purple indicating gaps in intron alignment when the intron is shared.

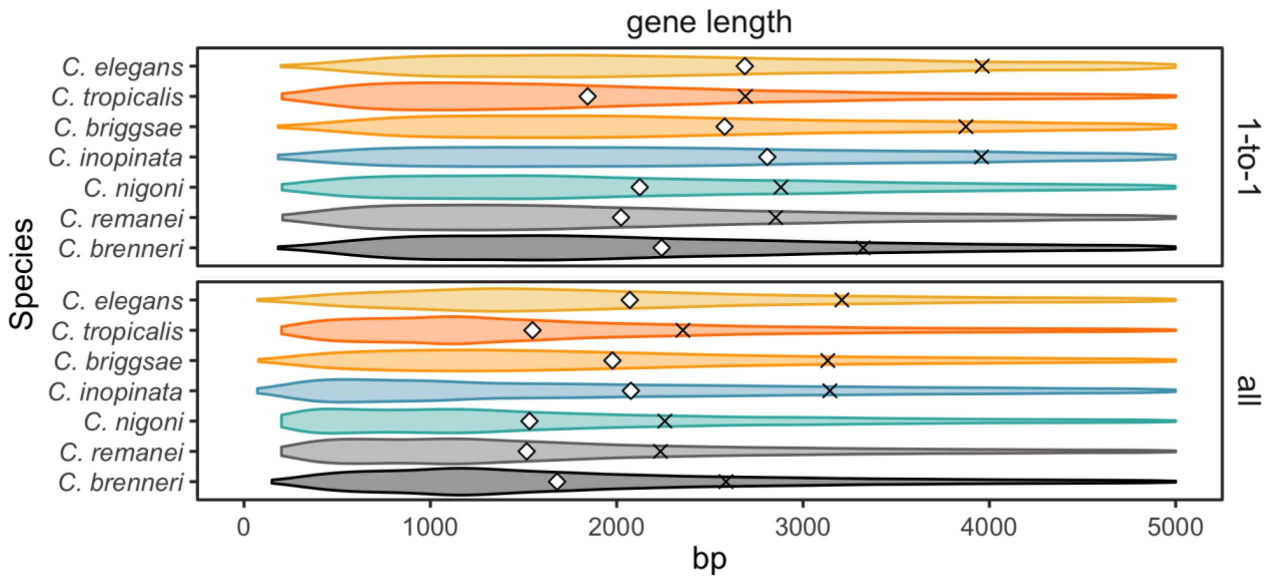


**Supplementary fig. S7**. Gene lengths in all genes and single-copy orthologs. The crosses represent the mean values, and the diamonds represent the medians.

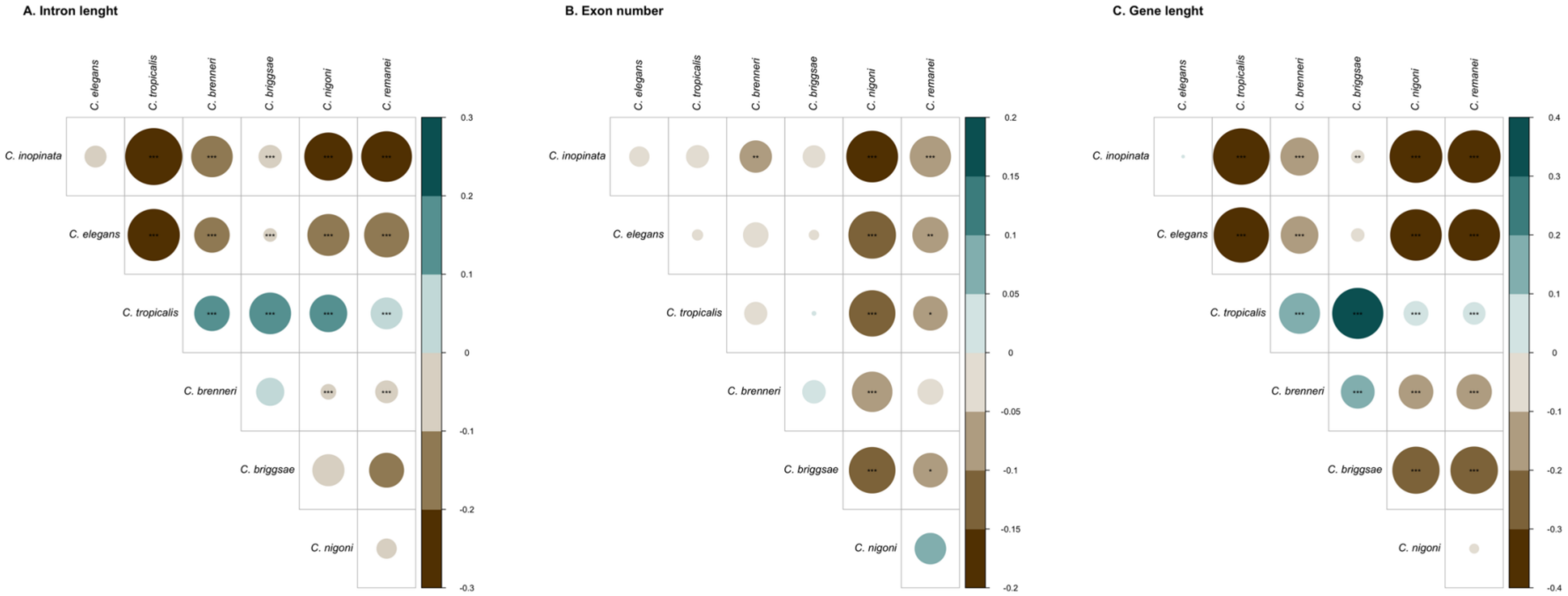


**Supplementary fig. S8**. Comparison of gene features in single-copy orthologs, where the size and color represent the value of Cohen's d. Asterisks indicate the Bonferroni-corrected p-values from non-paired Wilcoxon tests, where * represents adjusted p-value <0.1; ** represents adjusted p-value <0.001; *** represents adjusted p-value <0.0001. The specific comparisons illustrated are **(A)** intron lengths, **(B)** exon number, and **(C)** gene lengths.

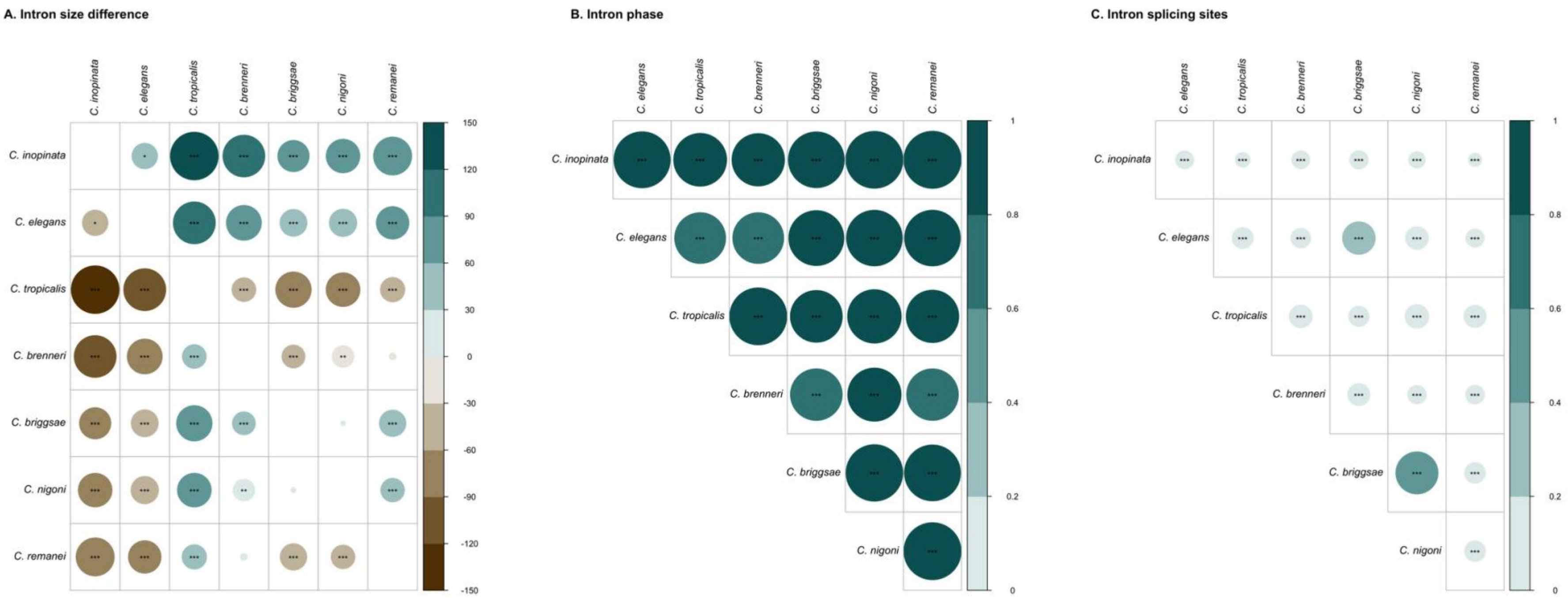


**Supplementary fig. S9 (A)** Comparison of corresponding intron sizes: The color and size of the circle indicate the mean difference in base pairs. Asterisks indicate the Bonferroni-corrected p-values from paired Wilcoxon tests, where * represents adjusted p-value <0.1; ** represents adjusted p-value <0.001; *** represents adjusted p-value <0.0001. Associations of **(B)** intron phase and **(C)** splicing sites and of corresponding introns in pairs of species: The color and size of the circles represent the value of Cramer's V. Asterisks indicate the Bonferroni-corrected p-values from chi-square tests, where * represents adjusted p-value <0.1; ** represents adjusted p-value <0.001; *** represents adjusted p-value <0.0001.
